# Host-dependent fungus-fungus competition suppresses fungal pathogenesis in *Arabidopsis thaliana*

**DOI:** 10.1101/2020.05.27.117978

**Authors:** Kuldanai Pathompitaknukul, Kei Hiruma, Hiroyuki Tanaka, Nanami Kawamura, Atsushi Toyoda, Takehiko Itoh, Yusuke Saijo

## Abstract

Like animals, plants accommodate a rich diversity of microbes, typically without discernible disease symptoms. How their pathogenesis is prevented in the host remains obscure. Here, we show that the root-infecting fungus *Colletotrichum fructicola* of the *C*. *gloeosporioides* clade (CgE), isolated from field-grown healthy Brassicaceae plants, inhibits growth of pathogenic fungi in *Arabidopsis thaliana*, in a phosphate status-dependent manner. Loss of host ethylene signaling or phytoalexins, camalexin or indole glucosinolates, however, allows CgE to display pathogenesis, suggesting host contributions to endophytic CgE colonization and benefit. Compared to a closely-related *C. gloeosporioides* pathogen (CgP), CgE is characterized by genome expansion and >700 fungal genes (4.34%) specifically induced in the host roots when co-inoculated with CgP, including genes related to fungal secondary metabolism. This may underlie antimicrobial tolerance of CgE and its dominance over pathogenic fungi within the host, pointing to a role for fungus-fungus competition in asymptomatic fungal colonization in plants.

## Introduction

In nature, plants are intimately associated with a rich diversity of microbial communities, including commensal, beneficial and pathogenic microorganisms (Bulgarelli *et al*., 2013; Lundberg *et al*., 2012; Duran *et al*., 2018; Toju *et al*., 2018). Plants often establish beneficial interactions with mutualistic microbes under adverse conditions. Most knowledge regarding mutualistic plant-microbe interactions has been obtained from several symbiosis models, including N_2_-fixing rhizobacteria and arbuscular mycorrhizal fungi, which promote host acquisition of nitrogen and phosphorus, respectively (Lugtenberg & Kamilova, 2009; Bonfante & Genre, 2010). Compared to these symbionts, despite their richness and diversity in root ecosystems, much less is known about the eco-physiological functions for fungal endophytes that colonize within living plants without causing diseases (Rodriguez, 2009).

Beneficial functions of fungal endophytes include plant protection from pathogens (Gao *et al*., 2010; Zhang *et al*., 2014). Direct protection relies on microbe–microbe competition between endophytes and pathogens, often with antifungal compounds (Zivkovic *et al*., 2010). *Trichodema harzianum* and *Serendipita vermifera* endophytes act as parasites to infect and suppress phytopathogens, thereby conferring host protection (Druzhinina *et al*., 2011; Moran-Diez *et al*., 2012; Sarkar *et al*., 2019). Species in *Trichoderma* genus also inhibit other fungi with antifungal secondary metabolites (Schuster & Schmoll, 2010). Many fungal toxins can be produced *in vitro* without microbial competitors or hosts (Gao *et al*., 2010; Kunzler, 2018), whereas a few of them specifically require microbial competitors (Konig *et al*., 2013). Conversely, some fungi employ ATP-binding cassettes and major facilitator superfamily transporters to detoxify or export fungal toxins (Morrissey & Osbourn, 1999; Gulshan & Moye-Rowley, 2007; Prasad & Goffeau, 2012; Ruocco *et al*., 2009). These attacking and defense mechanisms are likely to facilitate fungal competition with other microorganisms in the common host (Abdullah *et al*., 2017; Stroe *et al*., 2020). Whether and if so how host plants influence or exploit microbe–microbe competitions remain underexplored to date.

Beneficial bacteria and fungi also indirectly protect hosts by increasing local and/or systemic pathogen resistance (Van Wees *et al*., 2008; Pieterse *et al*., 2014; de Lamo & Takken, 2020). The endophytic ascomycete fungus *Harpophora oryzae* confers local and systemic rice resistance to rice blast fungi (*Magnaporthe oryzae*) (Xu *et al*., 2014). The basidiomycete *Serendipita indica* (formerly known as *Piriformospora indica*) induces systemic resistance in *Arabidopsis thaliana* against biotrophic powdery mildew, through the phytohormone jasmonic acid (JA) (Stein *et al*., 2008). However, molecular dissection of plant protection conferred by endophytic fungi has been hindered, in part due to the scarcity for genetic fungal studies in model plant species.

The ascomycete genus *Colletotrichum* causes anthracnose diseases in a wide range of crops, and is among the top 10 fungal pathogens of economic importance (Dean *et al*., 2012). Many *Colletotrichum* species are hemibiotrophic pathogens, displaying initial biotrophic and subsequent destructive necrotrophic phases (Perfect *et al*., 1999). In contrast to genuine obligate biotrophs such as powdery mildew and arbuscular mycorrhizal fungi, hemibiotrophic *Colletotrichum* species are amenable to axenic culture and genetic manipulation. In addition, high-quality genome sequences of over 10 species facilitate comparative genomics and molecular genetic studies in this genus (O’Connell *et al*., 2012; Gan *et al*., 2013 and 2016; Hacquard *et al*., 2016).

*Colletotrichum* genus has also endophytic species beneficial for the host plants. *C*. *tofieldiae* asymptomatically colonizes the roots of *Arabidopsis thaliana*, to promote plant growth under low-phosphate conditions. At the genome level, *Ct* is very closely related to root-infecting pathogenic species, such as *C. incanum* (Ci; Hacquard *et al.*, 2016). Indeed, even *Ct* displays high virulence in the host plants lacking tryptophan (Trp)-derived antimicrobial metabolites (Hiruma *et al.*, 2016). *Ct* overgrows and fails to promote plant growth in plants lacking MYB-type transcription factors *PHR1* and *PHL1*, two major regulators of phosphate starvation responses (PSR) (Hiruma *et al*., 2016). PSR enhances phosphate uptake and utilization under phosphate deficiency by reprogramming root system architecture and gene expression (Bustos *et al*., 2010), but how PSR serves to prevent fungal overgrowth remains obscure. High relatedness between beneficial and pathogenic species seems to be widespread, rather than exceptional, in plant-inhabiting fungi (Rodriguez *et al*., 2009). Pathogenic species/strains are often found, without displaying virulence, in microbial communities on apparently healthy plants (García *et al*., 2012; Xu *et al*., 2014). In *Arabidopsis thaliana*, root-inhabiting bacteria may contribute to asymptomatic accommodation of filamentous microbial eukaryotes, by antagonizing their negative impacts on the host (Duran *et al*., 2018). How potential virulence of pathogens or commensals is suppressed to achieve asymptomatic accommodation represents an important question in both plants and animals (Hiruma *et al*., 2018).

Here, we report an as-yet-undocumented beneficial *Colletotrichum* fungus, as well as its pathogenic relative, isolated from healthy field-grown cruciferous vegetables. Its colonization protects *Arabidopsis thaliana* plants from root-infecting fungal pathogens, in a manner dependent on ethylene, PSR and Trp-derived metabolites of the host. Transcriptional profiling in co-inoculated roots has produced an inventory of fungal genes that are specifically up- or down-regulated in the host-fungus-fungus interactions. Interestingly, both fungi strongly induce fungal genes related to fungal secondary metabolism. This implies chemical fungus-fungus competition dependent on the host, ultimately leading to suppression of fungal pathogenesis in plants.

## Results

### Isolation of endophytic and pathogenic *Colletotrichum* fungi from field-grown Brassicaceae vegetables

We have assembled a total of 116 fungal isolates from the asymptomatic roots and/or leaves of *Brassica* spp. after surface disinfection. Of them, we selected ten isolates for further analysis, based on the ease in cultivation and morphological and growth characteristics in culture, which were reminiscent of previously described fungal endophytes. We assessed inoculation effects of these fungi on *Arabidopsis thaliana* plants following fungal hypha inoculation in 1/2 × MS agarose media. Twenty-one d after individual inoculation, we detected varied effects among the tested strains, ranging from plant growth promotion to inhibition, indicated by shoot fresh weight (SFW) (Supplementary Fig. 1A and B). In particular, inoculation with fungal isolates E35, E41 and E66 increased SFW under nutrient-sufficient conditions, on average, by 89%, 51% and 115%, respectively, while in contrast E40 inoculation drastically reduced plant SFW by 68%, compared to mock controls. The results validate that healthy plants accommodate both plant growth-promoting (PGP) and pathogenic fungi.

Despite opposing effects on plant growth, pathogenic E40 and endophytic E41 fungi showed similar colony morphologies on potato dextrose agar (PDA) media, both characteristic of the *Colletotrichum* genus (Supplementary Fig. 1A). DNA sequencing of nuclear ribosomal internal transcribed spacer (ITS) regions, a universal DNA marker for fungal classification (Schoch *et al*., 2012), indicated that the two fungi were closely related to each other, within the clade *Colletotrichum gloeosporioides* (Supplementary Table 1), which we designated *C. gloeosporioides* pathogen (CgP) and *C*. *gloeosporioides* endophyte (CgE), respectively. Isolation of both fungi from apparently healthy plants prompted us to test the possible involvement of endophytes in suppression of CgP pathogenesis in the host.

### Endophytic CgE protects plants from pathogenic fungi

To examine what role CgE plays in host protection against pathogens, we co-inoculated CgP and CgE spores onto *Arabidopsis thaliana* roots. Inoculation of CgP alone resulted in severe inhibition of plant growth, indicated by a great decrease in SFW 21 d post inoculation (dpi) (Fig. 1A and B). By contrast, no discernible disease symptoms were observed when inoculated with CgE alone. Inoculation with CgE hyphae even promoted plant growth (Supplementary Fig.1). Importantly, CgE co-inoculation with CgP significantly reduced disease symptoms (Fig. 1A and B), compared with CgP inoculation alone. By contrast, co-inoculation of heat-killed CgE spores did not affect CgP infection (Fig.1B). These results indicate that live CgE fungi are required for host protection from CgP in *Arabidopsis thaliana*. We validated effectiveness of CgE-mediated protection against another root-infecting pathogenic species, *C*. *incanum* (Ci) (Sato *et al*., 2005; Hiruma *et al*., 2016; Hacquard *et al*., 2016), which is distantly related to CgP (Supplementary Fig. 2). These results suggest that CgE protection exceeds beyond niche competition within the *Colletotrichum gloeosporioides* species complex.

**Fig. 1.**
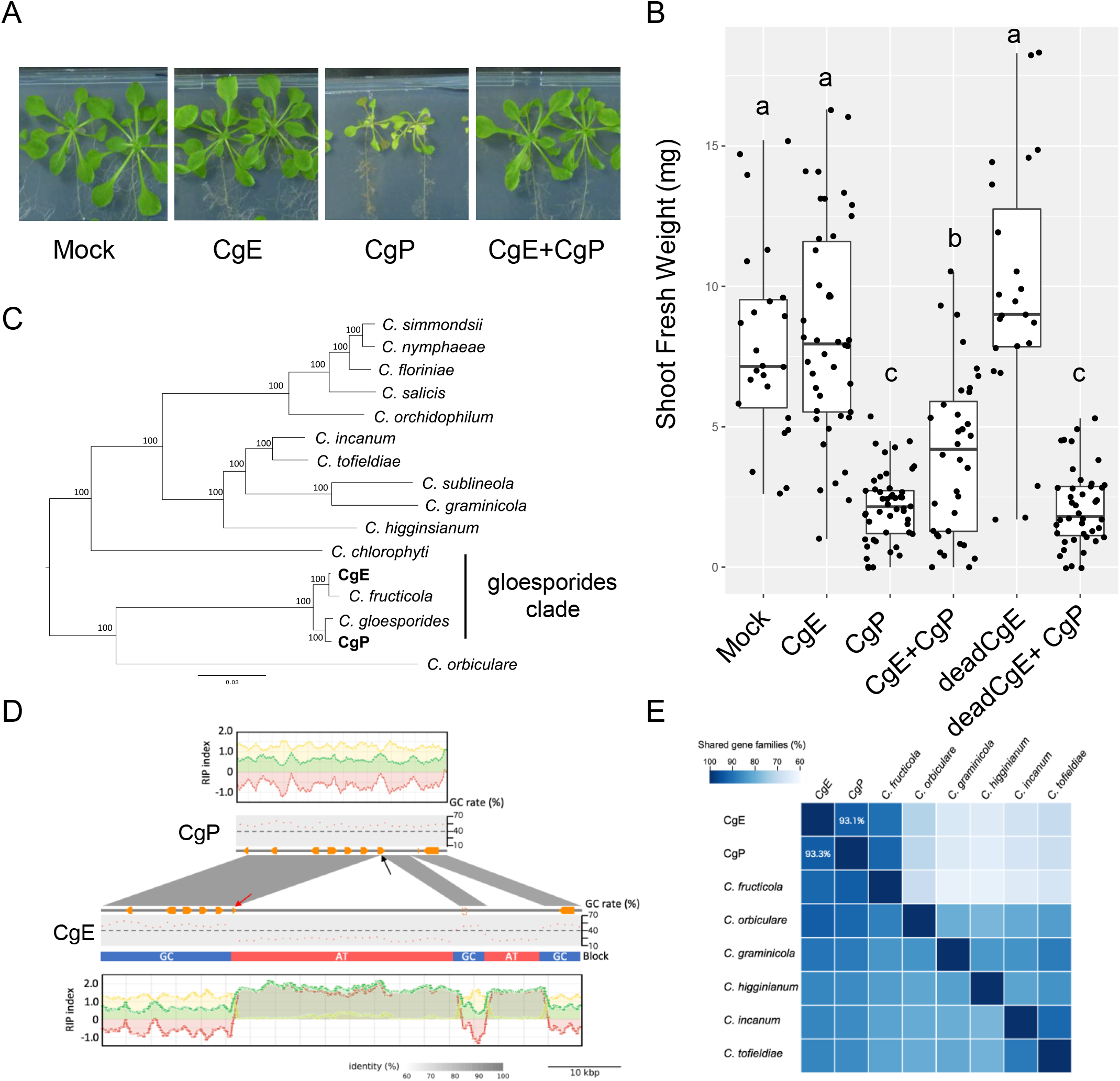
Endophytic CgE protects *Arabidopsis thaliana* plants from the closely-related pathogenic CgP. (A) Representative pictures of plants treated with water, endophytic *Colletotrichum* (CgE), and pathogenic *Colletotrichum* (CgP) or co-inoculated with CgE and CgP at 21 dpi on 1/2 MS agar media. (B) Shoot fresh weight of *Arabidopsis thaliana* from the co-inoculation assay at 21 dpi. Each sample comprised at least 10 shoots per experiment. The boxplot shows combined data from two independent experiments. The dots indicate individual replicates. Different letters indicate significantly different statistical groups (Tukey’s HSD, p < 0.05). (C) Phylogenetic positions of CgE and CgP. Phylogenetic tree of the concatenated protein-coding gene sequences for 16 *Colletotrichum* species. Multiple alignments of 4,650 ortholog groups were concatenated, and a phylogenetic tree was constructed by using 94,749 amino acid sites. Bootstrap values are shown on the branches. CgE and CgP belong to gloeosporioides clade. (D) One of the AT rich regions in CgE genome. AT block gene corresponds to *CGE00232* gene. The genomic sequences surrounding the *CGE00232* gene were extracted from the genome assembly of each allele in CgE and CgP. Vertical bars connecting adjacent genomic structures indicate BLAST hit blocks in the comparison between the two adjacent genomic scaffolds. Orange polygons indicate predicted genes. Red arrow indicates *CGE00232* gene. Sequences of *CGE00232* gene (Red arrow) and the dashed square regions show high similarity to *CGP01804* (Black arrow). RIP indexes of CgP (upper) and CgE (lower) are also described. RIP index values depicted: RIP product (green), RIP substrate (yellow) and RIP composite (red). RIP composite index values exceeding 0 indicate RIP activity. GC rate indicates GC content per 1 kb window. (E) Numbers of shared and specific orthologous family genes in CgE and CgP.

### CgE genome is characterized by long AT blocks with potential in generating SSP diversity

We obtained whole-genome information for CgE and CgP. Whole-genome alignment indicated CgE and CgP as different species in related taxa of the *Colletotrichum gloeosporioides* species complex (Fig. 1C). CgP genome size was approximately 57 Mb, similar to that of *C*. *fructicola* (previously described as *C*. *gloeosporioides*) Nara gc-5 strain (55.6 Mb, Gan *et al*., 2013), while CgE genome size was approximately 64.5 Mb, far larger compared to the other Cg strains sequenced to date (53.2–57.7 Mb) (Supplementary Fig. 3). Genome comparison revealed large AT-rich regions (GC content < 40%) as a unique feature of CgE genome, which largely explain increased genome size (Supplementary Fig. 4). Repeat‐induced point mutations (RIP) protect ascomycete fungal genomes against transposable elements, by converting C-G base pairs to T-A in duplicated sequences (Galagan & Selker, 2004). RIP indices (TpA/ApT dinucleotide ratios) were high in the AT-rich regions, in both genomes, (Fig. 1D), consistent with their generation by RIP. A specific feature of CgE, not conserved in CgP, included the existence of 13 genes predicted in AT-rich regions, which were all located near the borders with GC-rich regions (GC content > 40%). A border gene, *CGE00232*, appears to be generated via insertion of an AT-rich region into a conserved syntenic gene in CgP (Fig.1D). In GC-rich regions, both genomes had gaps at non-syntenic positions, at a considerably high frequency (Supplementary Fig. 6). Clustering protein-coding sequences into sets of orthologous genes with Proteinortho revealed that, of 15,763 CgE and 14,830 CgP gene families in total, 13,331 gene families were shared by the two fungi, while 2432 and 1499 gene families were specific to CgE and CgP, respectively (Fig.1E; Supplementary Table 2). These results indicate that the two genomes are more diverged than expected from the ITS sequences.

Fungal biotrophy relies on small secreted proteins (SSPs), which, if not all, contribute to suppression of host immunity (O’Connell *et al*., 2012; Lo Presti *et al*., 2015). The gene number of predicted SSPs was far the greatest in CgE among the sequenced Cg strains (Supplementary Table 2), despite similarity in the number of cell wall degrading enzymes (CWDEs), transporters, cytochrome P450 and secondary metabolite clusters (Supplementary Table 3, 4, 5, 6, and 7). This points to specific expansion of a SSP repertoire in CgE, in agreement with less destructive mode of infection.

### CgE inhibition of CgP growth is host dependent

We next tested whether CgE inhibition of pathogen growth occurs without the host, on a dual culture plate with CgE and CgP following inoculation onto the opposite sides. Although antibiotic Hygromycin B (100 μM) eventually suppressed colony growth of both fungi, CgE showed higher Hygromycin tolerance than CgP (Fig. 2A). By contrast, when co-cultured, CgP growth was not inhibited on the CgE side, suggesting that CgE does not directly inhibit CgP growth at least under the tested culture conditions.

**Fig. 2.**
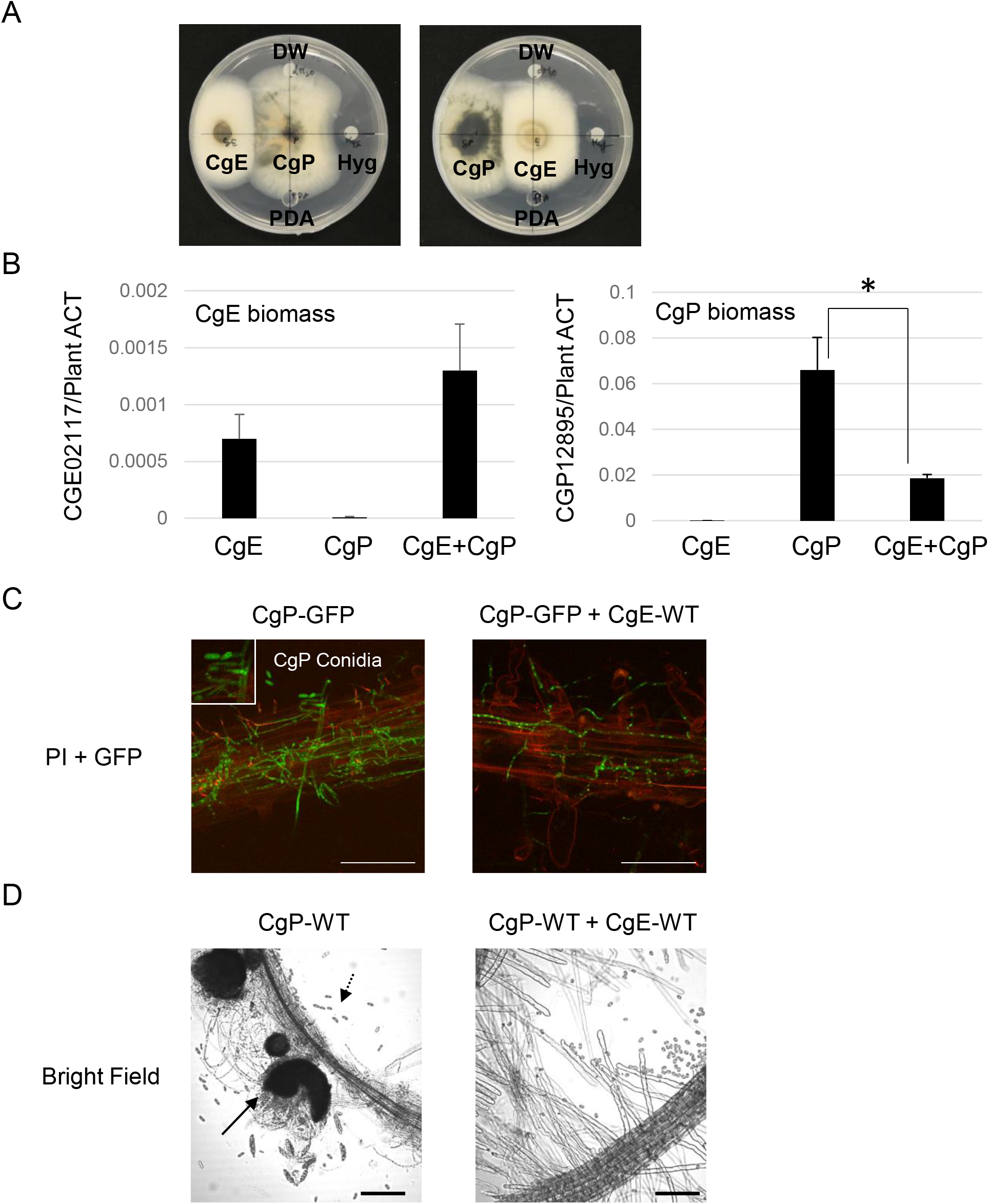
Endophytic CgE inhibits the growth of the pathogen CgP on *Arabidopsis thaliana* Roots. (A) Colony morphology of CgE and CgP at 7 dpi on PDA medium plates. Fungal colonies were placed beside another fungus, a PDA plug, and filter paper disc containing water or Hygormycin B. In contrast to Hygormycin B treatment, which formed an inhibition zone in front of CgP colony, CgE did not induce an inhibition zone in front of CgP. (B) A fungal biomass for CgE or CgP in roots was determined by qPCR using primers that specifically detect CgE or CgP genomes, respectively at 3 dpi. Bars represent means and standard deviation (SD) of data collected in 4 different root mixed samples (each sample comprised 20–25 roots) (\**t*-test, *p* < 0.01). (C) Confocal microscope images of CgP expressing cytoplasmic GFP (green) and *Arabidopsis thaliana* stained with propidium iodide (red). The hyphal network of CgP-GFP colonized around the root (left panel) and co-colonized with CgE-wild-type (Right panel). Arrows indicate the newly generated spores of CgP-GFP in the *Arabidopsis thaliana* root at 3 dpi. Scale bar, 100 μm. (D) Confocal microscope images of CgP and *Arabidopsis thaliana* plants. CgP formed spores (Left, dashed arrow) via black melanized structures (Left, black arrow). However, CgE inoculation inhibited formation of black melanized structures.

We then tested whether CgE restricts CgP growth *in-planta*, by quantitative PCR analysis with fungal species-specific primers (Fig.2B, Supplementary Table 13). CgP growth was greatly reduced 3 d post-inoculation (dpi) when co-inoculated with CgE, compared to CgP inoculation alone, while CgE growth was not affected by CgP (Fig. 2B). These results suggest that CgE outcompetes CgP in the host roots. We also employed transgenic CgP fungi constitutively expressing green fluorescence protein (CgP-GFP) under the control of *GPDA* regulatory DNA sequences from *Aspergillus nidulans* (O’Connell *et al*., 2007). Following co-inoculation of CgP-GFP with CgE, we traced live CgP growth and determined its abundance with the GFP signal as a proxy. Live imaging revealed that hyphal network of CgP-GFP in the roots was much less developed at 3 dpi in the presence of CgE than in its absence (Fig. 2C), suggesting that CgE restricts CgP hyphal growth at an early infection stage. In the absence of CgE, CgP produced new GFP-positive spores even at 3 dpi (Fig. 2C), and then formed numerous melanized structures at 10 dpi (Fig. 2D). These results suggest that CgE colonization inhibits growth and reproduction of CgP in *Arabidopsis thaliana* roots.

### Endophytic CgE colonization and host protection are phosphate status dependent

*C*. *tofieldiae* was reported to promote plant growth, specifically under phosphate deficiency in a manner dependent on the major PSR-regulating transcription factors *PHR1/PHL1* (Hiruma *et al*., 2016). In *phr1 phl1* plants, *C. tofieldiae* overgrows and fails to confer plant growth promotion, implying a role for PHR1/PHL1 in suppression of potential fungal pathogenesis. We tested possible phosphate status dependence of beneficial CgE interaction, in co-inoculation assays under normal (625 μM KH_2_PO_4_) and low phosphate (50 μM KH_2_PO_4_) conditions. CgP caused severe diseases irrespective of phosphate conditions (Figs 3A and B). Surprisingly, CgE also caused disease symptoms under low phosphate conditions, albeit to a lesser degree than CgP, and no longer protected the host despite slight alleviation of CgP pathogenesis (Figs 3A and B). These results suggest that CgE becomes pathogenic when phosphate is limited, in contrast to *C. tofieldiae*. Nevertheless, CgE disease symptoms became more severe in *phr1 phl1* plants, pointing to a critical role for *PHR1/PHL1* in restricting fungal pathogenesis under phosphate deficiency for both CgE and *C. tofieldiae* (Fig. 3C).

**Fig. 3.**
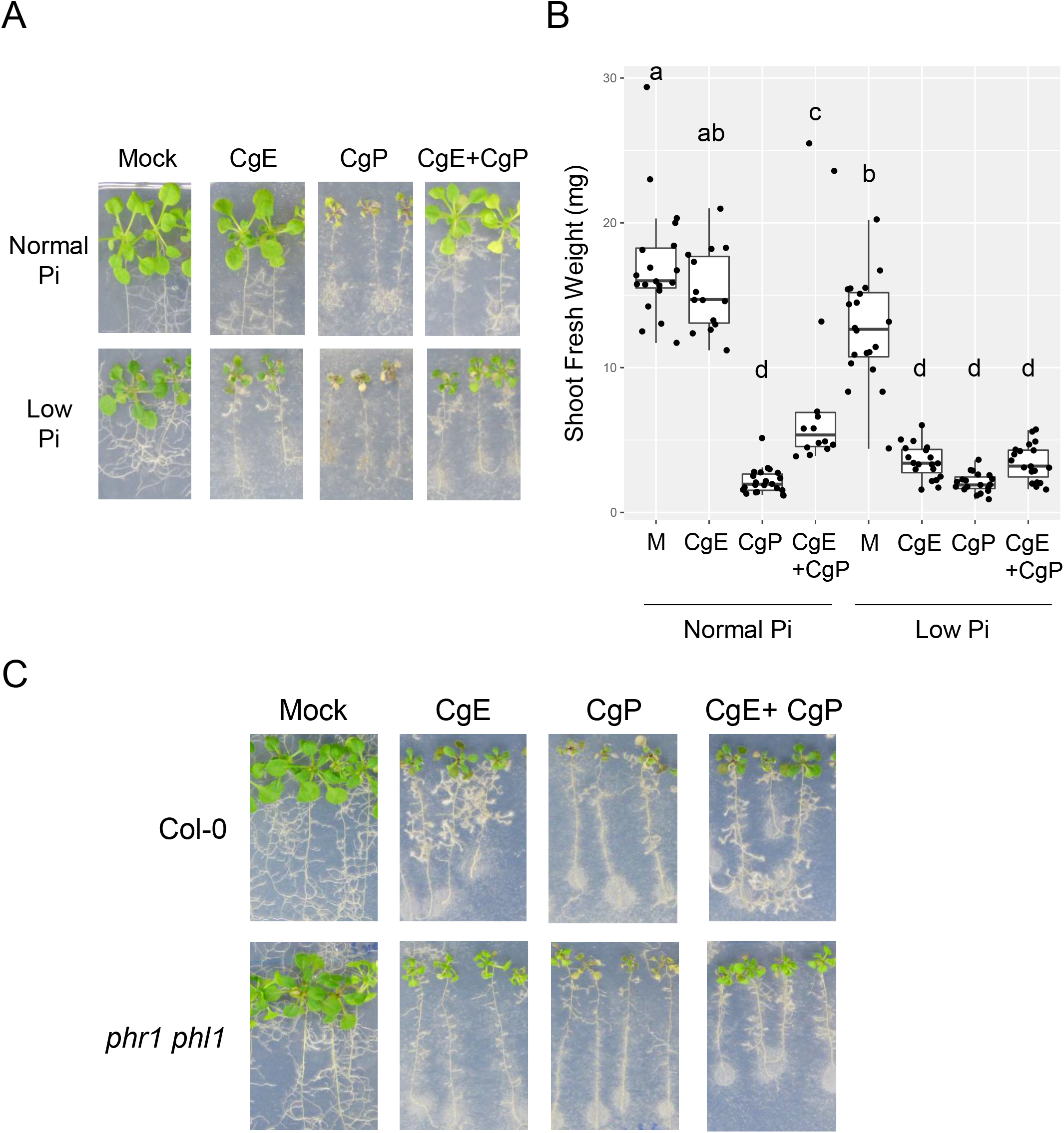
Endophytic CgE colonization and host-protective function are phosphate status dependent. (A) Morphology of plants treated with water, CgE, CgP or co-inoculation of CgE with CgP (CgE+CgP) at 21 dpi on 1/2 MS agar normal Pi (625 μM) and low Pi (50 μM) media. (B) Shoot fresh weight of *Arabidopsis thaliana* wild-type plants (Col-0) from the co-inoculation assay at 21 dpi. Similar results have been obtained in independent experiments. Different letters indicate significantly different statistical groups (Tukey’s HSD, p < 0.05). M= Mock. (C) Morphology of plants treated with water, CgE, CgP or co-inoculation of CgE with CgP on 1/2 MS Low Pi (50 μM) agar media.

### Endophytic CgE colonization and host protection require plant ethylene signaling

Ethylene, JA, and salicylic acid (SA) are among the major defense-related hormones that greatly influence plant-microbe interactions (Robert-Seilaniantz *et al*., 2011; Pieterse *et al.*, 2012). To determine the possible involvement of these hormone pathway(s) in beneficial interactions with CgE, we tested whether and if so how fungal infection modes and plant growth are influenced when the master regulator of ethylene signaling *EIN2*, enzymes required for JA and SA biosynthesis, *DDE2* and *SID2*, respectively, and SA signaling regulator *PAD4* are mutated. In *ein2 pad4 sid2* and *dde2 ein2 sid2* plants, CgE inoculation or co-inoculation with CgP resulted in severe growth retardation, pointing to a critical role for ethylene signaling in endophytic CgE colonization. By contrast, *dde2 pad4 sid2* plants largely retained WT-like growth and acquired CgP resistance after CgE inoculation (Supplementary Fig. 6A). These data suggest a pivotal role for host ethylene in the endophytic colonization and host-protective function of CgE.

We validated this notion in different ethylene-related mutants. *ein2-1* plants were hyper-susceptible to CgE, and were not protected by CgE against CgP (Fig. 4A and B). *ein3* and *eil1* plants, lacking ethylene-related transcription factors *EIN3* or *EIL1*, respectively (An *et al*., 2010), also displayed disease-like symptoms when inoculated with CgE (Fig. 4A and B, Supplementary Fig. 6B).

**Fig. 4.**
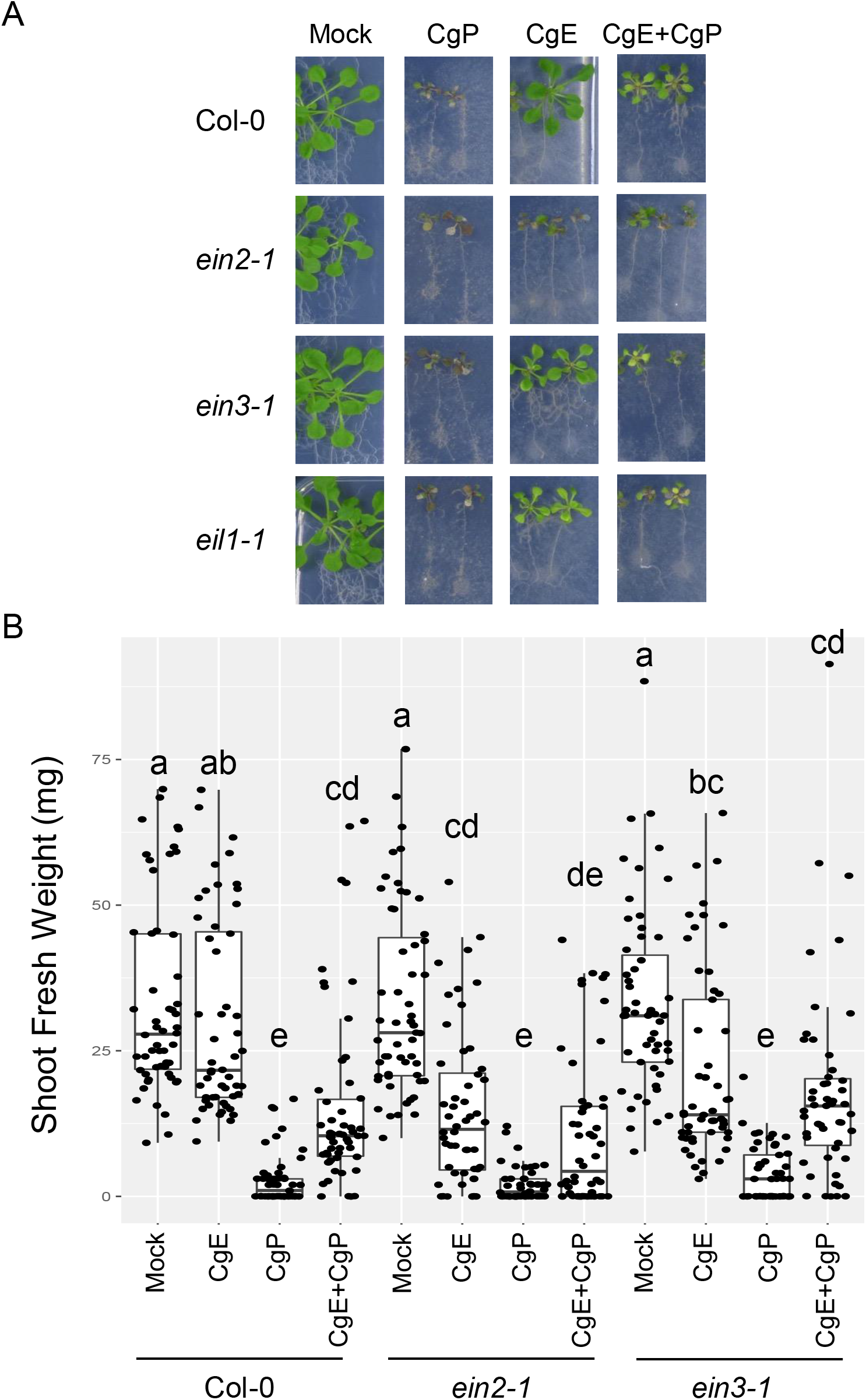
Endophytic CgE colonization and host protection require plant ethylene signaling. (A) Morphology of the plants treated with water, CgE, CgP or co-inoculation of CgE with CgP (CgE+CgP) at 21 dpi on 1/2 MS agar media. (B) Measurement of the shoot fresh weight of wild-type *Arabidopsis thaliana* and ET-related mutants (*ein2-1*, *ein3*) in the co-inoculation assay at 14 dpi. Each sample comprised around 20 shoots per experiment. The boxplot shows combined data from three independent experiments. Different letters indicate significantly different statistical groups (Tukey’s HSD, *p* < 0.05).

### Endophytic CgE colonization and host protection require host tryptophan (Trp)-derived metabolites

In *Arabidopsis thaliana*, Trp-derived secondary metabolites are required for the proper control of both pathogenic and endophytic fungi (Bednarek *et al*., 2009; Hiruma *et al*., 2016). As expected, loss of cytochrome P450-mediated conversion of Trp to indole-3-acetaldoxime, the initial catalytic step in this pathway (Fig. 5A), rendered *cyp79B2 cyp79B3* plants super-susceptible to CgP and also succumbed to CgE, allowing its pathogenesis (Fig. 5B).

**Fig. 5.**
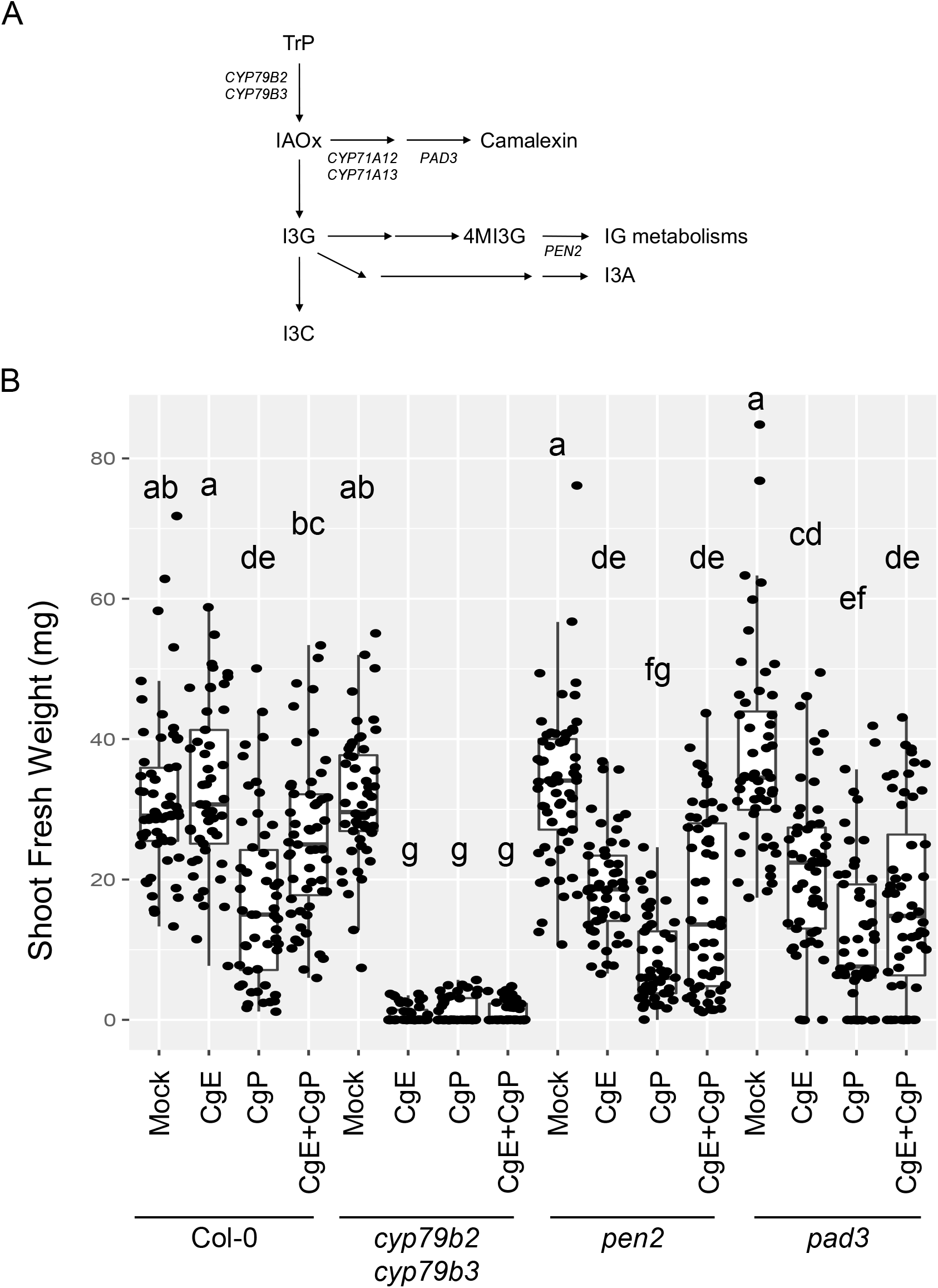
Endophytic CgE colonization and host protection require host tryptophan (Trp)-derived metabolites. (A) Scheme for Trp-derived metabolite pathways in *Arabidopsis thaliana*. (B) Measurement of the shoot fresh weight of *Arabidopsis thaliana* Trp-pathway mutant plants treated with water, CgE and CgP or co-inoculated with CgE and CgP at 14 dpi. The boxplot shows combined data from three independent experiments. Different letters indicate significantly different statistical groups (Tukey’s HSD, *p* < 0.05).

Disruption of *PENETRATION2 (PEN2)* atypical myrosinase (Lipka *et al*., 2005; Bednarek *et al*., 2009) or *PHYTOALEXIN DEFICIENT 3 (PAD3)* cytochrome P450 monooxygenase CYP71B5 required for antifungal camalexin biosynthesis (Zhou *et al*, 1999) also lost the control of CgE colonization and plant protection (Figs. 5A and 5B), as described for *C*. *tofieldiae* (Hiruma *et al*., 2016). As expected, *pen2* and *pad3* plants were both more susceptible to CgP than WT plants. These results indicate that potential virulence of CgE is de-repressed in the absence of host Trp-derived antimicrobial metabolites, and that its suppression is a key for beneficial interactions with CgE.

We then examined whether exogenous application of synthesized Trp-derived metabolites inhibits fungal growth in culture. In the presence of camalexin and indole-3-carbinol (I3C), growth of CgE and CgP was both suppressed, indicated by the colony diameters (Fig.5A, Supplementary Figs. 7A and 7B). By contrast, indole-3-ylmethylamine (I3A) did not suppress either growth (Supplementary Figs. 7A and 7B). These results suggest that specific subsets of Trp-derived antifungal metabolites (Camalexin, I3C and PEN2-dependent compounds excluding I3A, Fig. 5A) directly attenuate fungal growth to establish an endophytic mode in CgE. Interestingly, CgE again showed greater tolerance than CgP to camalexin and I3C in culture (Supplementary Figs. 7A and 7B), highlighting CgE tolerance to antifungal metabolites. This implies CgE adaptation to the root interior in *Arabidopsis thaliana*, wherein antifungal Trp-derived metabolites are highly induced in response to fungal challenge.

### Host transcriptome is not greatly altered during CgE colonization or protection

To have an overview of the host responses during CgE protection, we conducted RNA sequencing analysis in CgE-, CgP-, and co-inoculated roots at 6 h and 3 dpi. We first compared overall transcriptome profiles in multidimensional scaling analysis (Supplementary Fig. 8A). Although the host transcriptomes were not clearly separated between different inoculums and mock control at 6 hpi, fungal inoculation effects became apparent at 3 dpi (Supplementary Fig. 8A), implying intensive fungal challenge and/or host defense activation at this stage. Importantly, CgE and CgP inoculation differentially impacted the host transcriptome at 3 dpi (Supplementary Fig. 8A), consistent with striking differences in the host outcomes between the two fungi (Fig. 1A). Of particular note, root transcriptomes were nearly indistinguishable between CgE inoculation alone and co-inoculation with CgP, but far different from that of CgP inoculation alone, consistent with a collapse of CgP growth by CgE co-inoculation (Fig. 2B). CgE colonization essentially masked CgP effects on the host transcriptome.

Pairwise transcriptome comparisons [false discovery rate (FDR) < 0.01] revealed 13,300 differentially expressed genes (DEGs) at least in one of the pairs compared. These DEGs were classified into 12 different clusters by K-means clustering. Clusters 6, 11, and 12 were characterized by genes strongly responsive to both fungi, with Gene Ontologies (GOs) “response to chitin,” “innate immune response,” “Trp metabolism (tryptophan biosynthetic and metabolic process, glucosinolate biosynthetic and metabolic process),” and “plant hormonal response” dominating (Supplementary Fig. 8B, Supplementary Table 8). In clusters 11 and 12, in addition to defense responses, GOs related to hypoxia and ethylene signaling (ethylene-activated signaling pathway, response to ethylene, cellular response to ethylene stimulus) were overrepresented. Cluster 6, 11 and 12 were over-represented with genes strongly induced in response to CgP (Supplementary Fig. 8B, Supplementary Table 8), suggesting that CgP induces stronger defense activation than CgE, at this early interaction stage. Notably, this CgP effect was nearly abolished by CgE co-inoculation (Supplementary Fig. 8B), suggesting that host defense activation was alleviated in the presence of CgE.

### Fungal transcriptome dynamically changes during fungus–fungus competition in the host

CgE suppression of host transcriptional reprograming in response to CgP prompted us to examine fungal transcriptome during CgE-mediated host protection. We assembled *in-planta* fungal transcriptomes by separating CgE- and CgP-derived sequence reads in the co-inoculated roots, based on RNA-sequencing read mismatching to CgE and CgP genomes (Fig. 6A). Our method successfully identified the origin of the 93% sequence reads. Nearly a half of the total reads (48.13% ± 9.11%) were derived from CgE, whereas only a small portion (4.466% ± 0.32%) was derived from CgP (Supplementary Table 9). These results agree with CgE outcompeting over CgP (Fig. 2B and C).

**Fig. 6.**
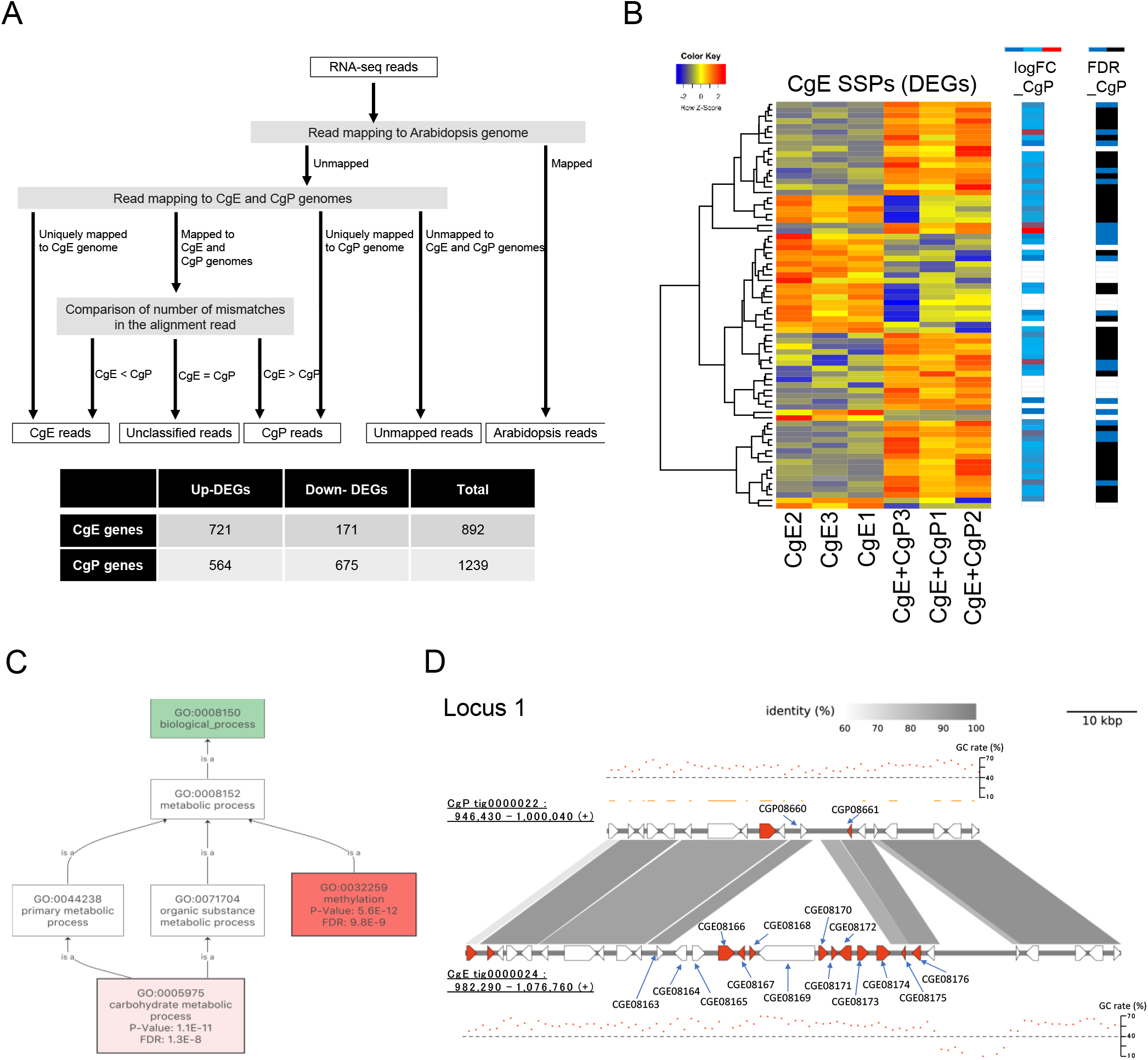
Transcriptome analysis of fungi detects large gene expression changes in co-inoculated samples. (A) Schematic diagram of the classification of RNA-seq reads from co-inoculated samples. The table represents a number of CgE or CgP genes specifically upregulated or downregulated in co-inoculated samples compared with that in the corresponding single-inoculated samples at 3 dpi. DEGs = differentially expressed genes. (B) Transcript profiling of 74 CgE SSP DEGs (|log_2_FC|⩾1, FDR<0.05) between CgE-colonized versus (vs) CgE+CgP-colonized roots. Overrepresented (yellow to red) and underrepresented transcripts (yellow to blue) are shown as log_10_ (read count +1). LogFC_CgP represents logFC (CgP-colonized vs CgE+CgP-colonized roots) of the corresponding CgP genes (Blue to Red). White represents the absence of obvious homologs in CgP (Similarity < 90%). FDR_CgP represents whether the expression levels of the corresponding CgP genes between CgP-colonized and CgE+CgP-colonized roots are significant (Blue: FDR < 0.05, Black: FDR > 0.05, White: no homologs in CgP (Similarity < 90%). (C) Results of Gene ontology (GO) analysis using 721 CgE up-regulated DEGs in the co-inoculated samples. The enriched GO terms of biological process were shown. (D) The expression profiles of CgE genes located in the secondary metabolism 25 cluster. The genomic sequences surrounding the secondary metabolism 25 cluster were extracted from the genome assembly of each allele in CgE and CgP. Vertical bars connecting adjacent genomic structures indicate BLAST hit blocks in the comparison between the two adjacent genomic scaffolds. Polygons indicate predicted genes. The red represents significantly higher expression in the co-inoculated samples compared to its alone (log_2_ FC >1, FDR < 0.05). GC rate indicates GC content per 1 kb window.

Next, we focused on *in-planta* fungal DEGs (|log_2_ FC| >1, FDR < 0.05) between individually-inoculated and co-inoculated roots. We detected 892 and 1,239 DEGs in CgE and CgP, respectively (Fig. 6A). Of the 721 CgE DEGs up-regulated in response to CgP, a large class were genes encoding SSPs, CWDEs (Supplementary note), secondary metabolite-related proteins including cytochrome P450, transporters, and antibiotic resistance proteins (Supplementary Table 10). For instance, of 659 CgE genes annotated for SSPs, 47 and 27 genes were up- and down-regulated following CgP co-inoculation, respectively (Fig. 6B). Although the majority of these CgE *SSP* genes (67 genes > 90%) were also conserved in CgP, most of CgP homologous genes (49 genes > 70%) displayed distinct expression patterns in the host (Fig. 6B). Of 618 *SSP* genes in CgP, 32 and 82 genes were up- and down-regulated, respectively, following CgE co-inoculation. Of these 114 CgP *SSP* genes, 104 genes were conserved in CgE genome but again displayed distinct expression patterns. These results suggest that the two fungi, despite close relatedness, express separate sets of SSPs during their competition in roots.

GO related to methylation (e.g., methyltransferases) dominated in CgE up-regulated genes following CgP co-inoculation (GO:0032259, FDR: 9.8E-9, Fig. 6C). Notably, 44 out of 94 *LaeA-like* (*LaeA* and *llm*) *methyltransferase* genes (annotated as Secondary metabolism regulator LAE1 or laeA) were highly induced in CgE during interaction with CgP in roots (log_2_ FC >1, FDR < 0.05, Supplementary Table 11). In different fungi, their homologues regulate production of secondary metabolites including fungal toxins (Palmer *et al*., 2013, Supplementary Fig 9). In addition, several methyltransferase genes other than LaeA-like were also highly induced during CgE-CgP competition. These data suggest the possible involvement of fungal secondary metabolites in the fungus-fungus competition. Indeed, CgE genes related to biosynthesis and efflux of secondary metabolites including fungal toxins, e.g., echinocandin B, T-2 toxin, botrydial, aspyridones, were over-represented in CgP-inducible DEGs, in the host (Supplementary Table 10). Furthermore, genes encoding cytochrome P450 monooxygenase, FAD-linked oxidoreductase, efflux pump, acyltransferase, prosolanapyrone synthase, C-factor, transcription factor and (N- and O-) methyltransferases in a CgE-specific secondary metabolism cluster (Cluster 25), closely located to an AT-rich region, were highly activated following CgP co-inoculation (Fig. 6D, Supplementary Fig. 5, Supplementary Table 10). Our results imply that CgE produces diverse secondary metabolites including fungal toxins in the host, in response to CgP, thereby suppressing CgP growth. Conversely, CgP also seems to produce different sets of fungal secondary metabolites in the host, in response to CgE, indicated by activation of some of *LaeA-like methyltransferase* genes (29 of 89 genes). Our results imply secondary metabolite-based fungus-fungus competition in the host (Supplementary Fig. 10; Supplementary Table 10 and 11).

Our *in vitro* culture assay revealed that CgP was more sensitive than CgE, to the antifungal compound echinocandin B, which inhibits synthesis of β-(1,3)-glucan (a major structural component of the fungal cell wall) (Walker *et al*., 2010, Supplementary Fig. 11), although the two fungi were essentially equally sensitive to another fungal toxin, aspyridone A (Macheleidt *et al*., 2016). Consistent with CgE tolerance to fungal toxins, an ABC transporter gene (*CGE02297*) related to fungal toxin efflux was strongly activated in CgE (Supplementary Table 10). The lack for dramatic transcriptome-wide changes in the host (Supplementary Fig. 8) implies that these toxins are specific to fungi (Walker *et al*., 2010). Our findings discover an important role for fungus–fungus competition, possibly via fungal toxins, in host protection by beneficial fungi.

## Discussion

### Contrasting lifestyles of two closely related *Colletotrichum* fungi in the host roots

In this study, we reveal two fungal species of the *C*. *gloeosporioides* clade, which is best known as devastating pathogens causing anthracnose diseases on important crops (Weir *et al*., 2012, Gan *et al*., 2013; Zhang *et al*., 2018). CgE is a root-associated endophyte conferring plant protection, whereas CgP is a highly virulent, root-infecting pathogen, both isolated from asymptomatic radish plants. Our results in *C*. *gloeosporioides* clade strengthen divergence of infection modes in *Colletotrichum* fungi, as described for *C. spaethianum* clade, including pathogenic (*C*. *incanum*) and beneficial (*C*. *tofieldiae*) species (Hiruma *et al*., 2016; Hacquard *et al*., 2016). These findings are consistent with the view that fungal pathogens have independently diversified infection modes in separate fungal lineages (Raffaele & Kamoun, 2012).

Increased genome size of CgE is largely explained by large AT-rich blocks. A similar case was reported between distantly related *C. orbiculare* 104-T and *C. fructicola* Nara gc5 (Fig.1A, Gan *et al*., 2013; 2019). Our evidence demonstrates genome expansion with AT-rich regions even within closely-related species of the same fungal clade. Such genome expansion is often associated with diversification of virulence factors such as SSPs or secondary metabolism-related genes in plant-infecting fungi (Rouxel *et al*., 2011). Of 13 CgE genes located in AT-rich blocks, expression of 5 genes has been validated during root colonization (Supplementary Table 12). Notably, genes located adjacent to an AT-rich block (secondary metabolism cluster 25) were highly induced specifically during fungal competition in the host (Fig. 6D), consistent with a role for AT-rich blocks in transcriptional activation (Nishi & Itoh, 1986; Palida *et al*., 1993). In *Epichloë* and *Neotyphodium* grass symbionts producing an extraordinarily diverse panel of anti-insect alkaloids, secondary metabolism clusters are present in proximity to AT-rich blocks (Schardl *et al*., 2013). AT-rich regions likely contribute to rapid evolution of part of microbial genomes, as illustrated in a “two-speed genome” model (Sanchez-Vallet *et al*., 2018), and stress-responsive gene regulation.

Comparative analysis for three different Cg genomes reveals that genes related to cell wall degradation, cytochrome P450 and secondary metabolite clusters are conserved in CgE and related pathogenic strains. Notably, however, *SSP* gene family is greatly expanded in CgE, of which some are specifically induced during competition with CgP in *Arabidopsis thaliana* roots (Fig. 6B, Supplementary Table 10). This is marked contrast to *SSP* repertoire in *C. tofieldiae*, which is reduced compared with closely related pathogenic species (Hacquard *et al*., 2016). Constraint of *SSP* repertoire in nonpathogenic relative to pathogenic species is also seen in *Fusarium oxysporum* (de Lamo *et al*., 2020). It is tempting to speculate that CgE utilizes SSPs to limit the opponent’s growth, rather than to promote infection, in the host.

### Endophytic colonization of *Colletotrichum fructicola* requires host ethylene

In *Arabidopsis thaliana*, CgE/CgP co-inoculation results in suppression of CgP virulence, consistent with their asymptomatic colonization in Brassicaceae vegetables. This requires ethylene-dependent suppression of potential CgE virulence. Ethylene signaling leads to production of pathogenesis-related proteins and phytoalexins as well as alterations in cell wall during pathogen resistance, in particular against necrotrophs (Thomma *et al*., 2001). Ethylene has also been implicated in beneficial interactions with mutualistic microbes (Zamioudis & Pieterse, 2012). *Fusarium oxysporum* disease suppression by endophytic *Fusarium solani* in tomato also requires the host ethylene (Kavroulakis *et al*., 2007). However, it is not clear whether ethylene signaling contributes to endophytic (non-pathogenic) colonization of *F. solani* by suppressing necrotrophy. Our findings with CgE extend a role for ethylene in suppression of fungal necrotrophy to endophytic species.

Remarkably, *C*. *tofieldiae*-mediated plant growth promotion is specific to phosphate deficiency, whereas CgE-mediated protection is specific to phosphate sufficiency (Fig.3). This provides compelling evidence for nutrition-dependent shifting of mutualistic fungal benefits and partners in plants. Interestingly, despite opposing effects of phosphate status on host benefits, both fungi overgrow in *phr1 phl1* plants under phosphate deficiency, pointing to a critical role for *PHR1/PHL1*-mediated PSR in restriction of fungal growth and pathogenesis (Fig. 3). PSR positively influences EIN3 protein accumulation during root hair formation under low phosphate conditions (Song *et al*., 2016; Liu *et al*., 2017). Conversely, *EIN3* and *EIL1* positively regulate *PHR1* expression in response to ethylene (Liu *et al*., 2017). Mutual positive feedback regulations between ethylene and PSR signaling may underlie *PHR1/PHL1*-mediated suppression of potential CgE pathogenesis. Ethylene also restricts biotrophic colonization of arbuscular mycorrhiza under low-Pi conditions (Kloppholz *et al*., 2011). However, *C*. *tofieldiae* colonization and plant growth promotion are unaffected by dysfunction of ethylene signaling (Hiruma *et al*., 2016), pointing to an ethylene-independent fungal control via PHR1/PHL1. *PHR1/PHL1* also negatively regulate SA-based defenses in assembling root-associated bacterial communities (Castrillo *et al*., 2017). How PHR1/PHL1 contribute to endophytic fungal colonization merits further in-depth studies.

### Fungus–fungus competition provides a basis for CgE-mediated host protection

Host-dependent CgE-CgP competition predicts the existence of a critical trigger for anti-fungal mechanisms in CgE, when it encounters a fungal competitor in the host. Host-dependent, competitor-induced extensive reprogramming of fungal transcriptome (Fig.6, Supplementary Fig. 9), without substantially affecting the host transcriptome (Supplementary Fig. 8), implies fungus-specific and –inducible nature of CgE competition mechanisms. Transcriptome data imply involvement of fungal secondary metabolites, including several toxins, during fungus-fungus competition. Fungal toxins are often produced as secondary metabolites under adverse conditions to the fungi, e.g. during host infection or anti-microbial defenses, and are often associated with fungal necrotrophy (Osbourn, 2010). Notably, however, CgE specifically induces these genes in the host, without impeding plant growth, in response to CgP.

Nearly a half (46%) of CgE *LaeA-like methyltransferase* genes are highly induced during competition with CgP in roots. LaeA is a putative methyltransferase that modulates heterochromatin structures (Bok & Keller, 2004; Reyes-Dominguez *et al*., 2010; Palmer *et al*., 2013). Deletion of *LaeA* in several fungal species lowers production of fungal secondary metabolites including fungal toxins, as well as fungal growth and virulence (Bok *et al*., 2006; Bouhired *et al*., 2007; Kale *et al*., 2008; Lodeiro *et al*., 2009; Wiemann *et al*., 2009). Conversely, CgP also induces different sets of fungal toxin-related genes and *LaeA-like methyltransferases* in the host, in response to CgE, albeit to a lesser degree compared with CgE. Increased numbers of *LaeA-like methyltransferase* genes in both CgE and CgP (94 and 86 genes, respectively) are notable compared to saprotrophic *Aspergillus nidulans* (10 genes) (Palmer *et al*., 2013). Repertoire expansion of fungal secondary metabolite regulators and their induction during host colonization in response to another fungus, suggests a critical role for fungal secondary metabolites in fungus-fungus competition. Acquisition of a fungal toxin-detoxifying enzyme gene of endophyte origin in wheat confers *Fusarium* head blight resistance (Wang *et al*., 2020). In bacteria, type IV and VI secreted systems are employed to directly inject toxins to eukaryotic and bacterial competitors (Basler *et al*., 2013; Ma *et al*., 2014; Souza *et al*., 2015; Trunk *et al*., 2018; Kim *et al*., 2019). Interestingly, *Pseudomonas aeruginosa* appears to sense the presence of functional type VI secretion in *Vibrio cholerae* competitor (Basler *et al*., 2013). These studies and ours suggest the existence of mechanisms by which infectious microbes respond to competitors in the host environment. How the host influences microbe-microbe competitions for host benefits merits future studies.

## Materials and methods

### Plant–fungus interaction assay by plant and fungal cocultures

CgE and CgP were mainly used for plant-fungus interaction assay. In brief, 7-day-old plants grown on 1/2 MS media with 25 mM sucrose were placed to 1/2 MS media without sucrose in 9-cm square plates. Spore suspensions of CgE, CgP and the mixed suspension were dropped onto the plant root tips (5 μl each plant). The initial spore suspension of CgE and CgP in each treatment was adjusted to the same amount (25 spores/plant). The mixed suspension contained the same amount of CgE and CgP spores (each 25 spores/plant). Dead spores were prepared by autoclaving (121°C, 15 min). Plates were placed horizontally in a temperature-controlled room with a photoperiod of 12-h light/12-h dark and temperature of 21°C ± 1°C. Full details are given in Supplementary Experimental Procedures.

### Fluorescence microscopy

Inoculated *Arabidopsis thaliana* roots were visualized using fluorescence microscopy. The studies were performed using a confocal laser scanning microscope Olympus FV1000 with excitation at 488 nm for GFP or bright field and 560 nm for propidium iodide (PI) at 10×, 20×, and 40× magnification. PI (10 mg/ml) was used to stain root cell walls by direct application onto the slide.

### Genome sequencing and assembly

Fungal DNA was extracted by CTAB with RNase treatment from fungal hyphae grown on liquid Mathur’s medium (glucose 2.8 g/l, MgSO_4ֺ7_H_2_O 1.2 g/l, KH_2_PO_4_ 2.7 g/l, and mycological peptone LP0040 2.2 g/l) for 2 days. Genomic sequences of CgE and CgP were determined using PacBio single-molecule real-time sequencing and Illumina HiSeq for paired-end short reads. Genome sequences for CgE and CgP are deposited in DDBJ (DRA009690, CgE=CfE). Full details about genome sequencing, genome assembly, gene prediction, gene annotation and comparative genomics are given in Supplemental Experimental Procedures.

### Transcriptome analysis

RNA samples were extracted from inoculated roots at post-inoculation 6 h and 3 dpi. Total RNA was extracted using a NucleoSpin RNA Plant (Macherey-Nagel). RNA samples (1μg each) were then sent to Macrogen for library preparation and subsequent sequencing. RNA-seq read sets obtained from CgE, CgP, and co-inoculated samples were subjected to adapter removal and quality filtering using Platanus_trim (version 1.0.7) with default parameters. The trimmed reads were classified using two sequential rounds of mapping. First, the trimmed reads were mapped to the *Arabidopsis thaliana* genome using HISAT (version 2.1.0) (Kim *et al*., 2015) with default parameters. Reads that were mapped onto the *Arabidopsis thaliana* genome were classified as originating from *Arabidopsis thaliana*. Next, we performed the second classification by mapping the reads that remained unmapped in the first classification onto CgE and CgP genomes. Reads that uniquely mapped to either CgE or CgP genomes were classified as originating from CgE or CgP, respectively. In addition, reads that could not be mapped to both genomes were classified as unmapped reads. Finally, reads that mapped to both genomes during the second step were further classified. The number of mismatches in the alignment reads were identified by the “XM” flag in the SAM output files and then the number of mismatches in the alignment against CgE and CgP genomes were compared. Based on the comparison results, reads were classified into CgE, CgP, and classified reads. The resulting CgE-RNA-seq derived CgE reads (CgE-CgE reads), CgP-RNA-seq derived CgP reads (CgP-CgP reads), co-inoculation-RNA-seq derived CgE reads (coinoc-CgE reads), and co-inoculation-RNA-seq derived CgP reads (coinoc-CgP reads) were used for differential expression analysis. RNAseq sequences used in this study are deposited in DDBJ (DRA009854, CgE=CfE). Full details are given in Supplementary Experimental Procedures.

Supplementary information includes Supplementary Experimental Procedures, Supplementary note, 11 figures, and 14 table.

## Contributions

KH, YS conceived the study. PK, KH, HT, NK, and AT conducted the experiments. PK, KH, HT, NK, AT and TI analyzed the data. PK, KH, and YS wrote the paper with feedback from all the co-authors.

## Acknowledgements

We thank Ms. Mie Matsubara, and Ms. Akemi Uchiyama for technical assistance, and Dr. Shigetaka Yasuda for advice on fungal growth assays with tryptophan-derived metabolites. We thank Dr. Yasuyuki Kubo for Agrobacterium C58C1 strains. PK was supported from The NAIST International Scholarship. This work was supported in part by Japan Society for the Promotion of Sciences (JSPS) KAKENHI Grant (16H06279, 16KT0031, 18H04822, 18K14466, 18H02467), the Japan Science and Technology Agency (JST) grant (JPMJPR16Q7, JPMJSC1702), and Asashi Grass Foundation. Computations were partially performed on the NIG supercomputer at ROIS National Institute of Genetics.

