## Supplementary information and Figures for "Host-dependent fungus-fungus competition suppresses fungal pathogenesis in *Arabidopsis thaliana*"

#### Supplemental information

Supplementary information includes Supplementary Experimental Procedures, Supplementary note, 11 figures, and 14 table.

#### Materials and methods

##### Fungal isolation

Daikon radish (925009, SaKaTa seeds) were grown in an open field (Green Lab, Nara Institute of Science and Technology, GPS: 34.734615, 135.736572) with conventional chemical fertilizer (N:P:K = 8:8:8 (LIFELEX), 80g /$m^2$) before planting from November 2015 to January 2016. Mature healthy plants were harvested and washed for 15 min under running tap water; shaken for 1 min in 70% ethanol followed by 30 s in 1% sodium hypochlorite; and finally washed three times in sterilized water. Plant parts were left to dry and cut into small pieces of approximately 3 × 3 mm and then placed on BD Difco™ PDA (Becton, Dickinson and Company) in 60-mm petri dishes. After 3 days, pure cultures were obtained by picking hyphal tips of visible colonies growing out of plant parts. Hyphae were transferred to new PDA media and incubated at 25°C ± 1°C. Fungal isolates were also cryopreserved at -80°C as 25% glycerol stocks.

##### Molecular identification of endophytic fungi

The ITS region on the ribosomal DNA was amplified using the primers ITS1 and ITS4 (Supplementary Table 13). PCR was performed using KOD FX Neo® (Toyobo Life Science). PCR products were purified using a FastGene Gel/PCR Extraction Kit (Nippon Genetics). These samples were subjected to cycle sequencing using a BigDye™Terminator Cycle Sequencing reaction in a PCR machine. The sequencing samples were then purified by ethanol/EDTA precipitation. The purified samples were sequenced using an ABI 3100 sequencer. Sequences were assembled and edited using BioEdit and then submitted to a BLAST search.

##### Plant material and growth conditions

*Arabidopsis thaliana* accession Col-0 and the *ein2 pad4 sid2* (Tsuda *et al.*, 2009), *ein2 sid2 dde2* (Tsuda *et al.*, 2009), *ein2 dde2 pad4* (Tsuda *et al.*, 2009), *sid2 pad4 dde2* (Tsuda *et al.*, 2009), *ein2-1* (Alonso *et al.*, 1999*)*, *ein3* (An *et al.*, 2010), *eil1* (An *et al.*, 2010), *cyp79B2 cyp79B3* (Zhao *et al*., 2002*)*, *pen2 (*Bednarek *et al*., 2009) , *pad3* (Zhou *et al*., 1999) and *phr1 phl1* (Bustos *et al*., 2010) mutants were used for the study of plant interactions with endophytes and pathogens. Seeds were surface sterilized with 70% ethanol for 30s followed by 6% sodium hypochlorite with 0.01% Triton X for 15 min. After being washed three times in sterilized water, seeds were placed in cold treatment at 4°C for 72 h. The treated seeds were sown on 1/2 MS media with 25 mM sucrose. Plates were placed vertically in a plant growth chamber with a photoperiod of 12-h light/ 12-h dark at a temperature of 21°C ± 1°C. For the inoculation assay, plants were transferred to 1/2 MS media without sucrose.

**Fungal screening by plant and fungal cocultures**

Fungi with unique colony morphology and/or spore formation in our media were selected for the further assay. To avoid contamination, we did not test fungi that could spread their spores easily by air. To observe their interactions with the host, two agar plugs (7 mm diameter) from 10-day-old fungal samples were cut with a sterile cork-borer and transferred to 1/2 MS media in 9 cm square plates. After 5 days, 7-day-old plants grown in half MS media with 25 mM sucrose were placed at a distance of 2 cm from the edge of the fungal colonies. A sterile PDA plug was used as negative control. Plates were placed horizontally in a temperature-controlled room with a photoperiod of 12-h light/12-h dark (~ 40 μ mol m-2 s-1) at a temperature of 21°C ± 1°C. Unless otherwise mentioned, plant SFW was measured 21 days after infection.

**Dual culture assay *in vitro***

Agar plugs of 10-day-old CgP and CgE cultures were transferred to PDA plates; the agar plugs were placed 2.5 cm apart (edge-to-edge). Sterile paper discs containing 5 μl of 10 mM of Hygromycin B were used as positive controls. Agar plugs for non-cultured PDA plates and paper discs containing 5 μl of sterile water were used as negative controls. Colony formation and inhibition rates were determined 10 days after inoculation by measuring the colony radius from center-to-edge.

**Assay of fungal inhibition by Trp-derived secondary metabolites and fungal toxins**

Agar plugs of 10-day-old cultures of CgP and CgE were transferred to half strength MS media containing 100 μM Trp-mediated metabolites (Camalexin (SIGMA-ALDRICH (#SML1016)), I3A (Nacalai (#54588-31)), and I3C (Tokyo chemical industry (#I0496))) in 60 mm petri dishes.

Medium containing dimethyl sulfoxide (DMSO) was used as a negative control. Colony formation and inhibition were observed 10 days after inoculation by measuring the colony radius from center-to-edge. For fungal toxin assay, cultures of CgP and CgE were transferred to PDA media supplemented with or without fungal toxin echinocandin B (4 µg/ml, E) or aspyridone A (4 µg/ml, A). Colony formation and inhibition were observed 5 days after inoculation.

**Plant–fungus interaction assay by plant and fungal cocultures**

CgE, CgP or *C. incanum* were used for plant-fungus interaction assay. In brief, 7-day-old plants grown on 1/2 MS media with 25 mM sucrose were placed to 1/2 MS media without sucrose in 9-cm square plates. Recipe of half MS low Pi medium used in this study are described in Gruber et al. Spore suspensions of CgE, CgP and the mixed suspension were dropped onto the plant root tips (5 µl each plant). The initial spore suspension of CgE and CgP in each treatment was adjusted to the same amount (25 spores/plant). The mixed suspension contained the same amount of CgE and CgP spores (each 25 spores/plant). Dead spores were prepared by autoclaving (121°C, 15 min). Plates were placed horizontally in a temperature-controlled room with a photoperiod of 12-h light/12-h dark and temperature of 21°C ± 1°C. Plant SFW and plant morphology were observed at 14 or 21 days after infection.

**Quantitative PCR analyses of fungal biomass**

Genomic DNAs were extracted from *Arabidopsis thaliana* roots inoculated with fungi by CTAB method (2% cetyl trimethylammonium bromide, 1% polyvinyl pyrrolidone,100 mM Tris-HCl, 1.4 M NaCl, and 20 mM EDTA). 30 ng genomic DNAs were used for qPCR analysis using SYBR™ green (Applied Biosystems) on a Thermal Cycler Dice® Real Time System II (Takara). *ACTIN2* (At3g18780) were used as internal controls and the relative expression level of each gene was calculated by the $2^{-\Delta\Delta Ct}$ method. For qRT-PCR analysis, total RNAs were extracted using the Purelink™ RNA Purification solution (Invitrogen), followed by chloroform purification and alcohol precipitation. A 500-ng quantity of the RNAs was reversely transcribed to cDNA using Primescript™ reverse transcriptase (Takara).

**Fungal transformation via *Agrobacterium*-mediated transformation**

The constitutively expressing GFP lines of CgP were generated by *Agrobacterium*-mediated transformation using the binary plasmid pBin-GFP-hph (O'Connell *et al.*, 2004) with the GFP gene under the control of the constitutive GPDA promoter. Fungal transformants were selected by PDA containing 100 μM Hygromycin B.

**Fluorescence microscopy**

Inoculated *Arabidopsis thaliana* roots were visualized using fluorescence microscopy. The studies were performed using a confocal laser scanning microscope Olympus FV1000 with excitation at 488 nm for GFP or bright field and 560 nm for propidium iodide (PI) at 10x, 20x, and 40x magnification. PI (10 mg/ml) was used to stain root cell walls by direct application onto the slide.

**Genome sequencing and assembly**

Fungal DNA was extracted by CTAB with RNase treatment from fungal hyphae grown on liquid Mathur's medium (glucose 2.8 g/l, $MgSO_4{\cdot}7H_2O$ 1.2 g/l, $KH_2PO_4$ 2.7 g/l, and mycological peptone LP0040 2.2 g/l) for 2 days. Genomic sequences of CgE and CgP were determined using PacBio single-molecule real-time sequencing and Illumina HiSeq for paired-end short reads. For PacBio sequencing, a SMRTbell library was constructed using the SMRTbell Express Template Kit (Pacific Biosciences, CA, USA) according to the manufacturer's protocol. The sequencing library was size-selected using the BluePippin system (Saga Science, MA, USA) with a minimum fragment length cutoff of 30 kbp. One SMRT cell (1M v2) was run on the PacBio Sequel System with Binding Kit2.0/Sequencing Kit2.1 and 600 min movies. In addition, genomic DNA was fragmented to an average size of 600 bp with the DNA Shearing System M220 (Covaris Inc., MA. USA). A paired-end library was constructed with a TruSeq DNA PCR-Free Library Prep kit (Illumina, CA, USA) and was size-selected on an agarose gel using a Zymoclean Large Fragment DNA Recovery Kit (Zymo Research, CA. USA). The final library was sequenced on the Illumina HiSeq 2500 sequencer with a read length of 250 bp. The PacBio sequence reads were trimmed and assembled initially using Canu (version 1.6) (Koren *et al.*, 2017). This initial assembly was polished to correct sequencing errors twice. The first polishing event was performed using Arrow (version 2.2.1) (Chin *et al*. 2013) with PacBio long reads alignment from PBalign (version 0.3.1). The second polishing event was performed using Pilon (version 1.22) (Walker *et al*. 2014) with Illumina short reads alignment from BWA (version 0.7.15) (Li & Durbin 2010). Genome sequences for CgE and CgP are deposited in DDBJ (DRA009690, CgE=CfE).

**Comparison of the genomic sequences between CgE and CgP**

For the whole-genome comparisons, assembled scaffold sequences longer than 100 kbp in length were used. Pairwise-alignment was performed using "nucmer" (MUMmer version 3.1) program. Alignment results were filtered using the "delta-filter" (MUMmer version 3.1) program with the '-l 5000' parameter. The result of dot-plot analysis between CgE and CgP genomes was visualized using "mummerplot" (MUMmer version 3.1).

For the compilation of the genomic region surrounding CgE-specific genes, pairwise-alignment was performed using blastn algorithm, then blast HSPs (High-scoring Segment Pairs) less than 1,000 bp were filtered. The identity and structural differences of the corresponding genomic region were visualized based on blast results using an in-house Python script. RIP analysis was performed using the RIPper (Van Wyk *et al.*, 2019). RIP indices were calculated for all sequences, using a 1,000 bp sliding window and 200 bp step.

**Gene prediction and annotation**

Protein-coding region identification and gene prediction were conducted by a combination of RNA-seq-based prediction, homology-based prediction, and *de novo* prediction methods. For RNA-seq-based prediction, RNA-seq reads were mapped to assembled genomes using HISAT (version 2.1.0) (Kim *et al.* 2015) with default parameters to identify exon regions and splice positions. Gene structure predictions were made by StringTie (version 1.3.4d) (Pertea *et al.*, 2015) with default parameters. For homology-based prediction, protein sequences from 14 species (*C. fructicola* strain Nara gc5, *C. gloeosporioides* strain cg14, *C. graminicola* strain M1.001, *C. higginsianum*, *C. orbicular*, *C. tofieldiae*, *C. chlorophyte*, *C. fioriniae*, *C. incanum*, *C. nymphaea*, *C. orchidophilum*, *C. salicis*, *C. simmondsii*, and *C. sublineola*) were aligned to assembled genomes using Exonerate (version 2.2.0) (Slater & Birney, 2005). For ab initio gene prediction, GeneMark-ES (Version 4.33) and BRAKER (Version 2.1.0) were used (Ter-Hovhannisyan *et al.*, 2008). GeneMark-ES was trained on its own genome sequence and the resulting bam file from HiSat2 was used to train BRAKER. All gene models predicted from the above three approaches were combined by EvidenceModeler (version 2.2.0) into a nonredundant set of gene structures (Haas *et al.*, 2008).

For functional annotation, the predicted gene sequences were searched with various databases such as NCBI nt, Swiss-Prot (Bairoch & Boeckmann, 1991), and InterProScan (Jones *et al.*, 2014), with a cut off E value of $1E\text{-}10^{5}$. The prediction of subcellular localizations was obtained by SignalP (Nielsen *et al.*, 1997). For the prediction of secreted proteins, we identify N-terminal signal peptides and exclude sequences with transmembrane domains and GPI-anchors by TMHMM (Krogh *et al.*, 2001) and Fungal BIG-PI (Eisenhaber *et al.*, 2004). The sequences were then submitted to TargetP (Emanuelsson *et al.*, 2000) analysis to identify predicted sequences for extracellular localization. Potential secondary metabolite clusters and associated backbone genes were identified using SMURF (Khaldi *et al.*, 2010). Carbohydrate active enzymes were classified using the dbCAN HMMER - based classification system (Yin *et al.*, 2012) applying an E - value cut - off of 1E-$10^5$. Transporters were annotated by BLAST search of the Transporter Classification Database (Saier *et al.*, 2009) using BLASTp (E-value cut-off 1E-$10^5$, similarity cut - off 30%, alignment coverage > 50%). Cytochrome P450 proteins were first identified as proteins with homology in Fungal cytochrome P450 database (Park *et al.*, 2008) using BLASTp (E-value cut-off 1E-$10^5$, similarity cut - off 30%). Among them, proteins with PF00067 domain were finally determined as cytochrome P450 proteins. Candidates of LaeA-Like proteins were first identified as proteins with homology to LaeA proteins in Swiss-Prot (E-value cut-off 1E-$10^5$, alignment coverage > 70%). Among them, proteins with S-adenosylmethionine-dependent methyltransferase domain (IPR029063, cd02440) were finally determined as LaeA-Like proteins.

**Construction of a phylogenetic tree of concatenated protein-coding gene sequences**

We determined the orthologous groups of proteins from 16 *Colletotrichum* species (see the "Orthologs among *Colletotrichum* species" section) and extracted 4,650 groups with a one-to-one relationship across all species. For each group, multiple alignments were performed by MAFFT (Katoh & Standley 2013) and sites containing gaps ("-") or ambiguous characters ("X") were excluded. All alignments were concatenated, and 94,749 amino acid sites were used for phylogenetic analysis. A phylogenetic tree was constructed with RAxML (version 8.2.12) (Stamatakis 2014). Here, we applied the JTT substitution matrix with a gamma model of rate heterogeneity (-m PROTGAMMAJTT) and 1000 replicates for bootstrap analysis.

**Orthologs among *Colletotrichum* species**

We classified all proteins from 16 *Colletotrichum* species into orthologous groups using Proteinortho (Supplementary Table 13, Lechner *et al.*, 2011). This tool constructs groups on the basis of an all-against-all alignment of BLASTP. The definition of an orthologous relationship between proteins was as follows: e value ≤ 10 to 5; identity ≥ 25%; and alignment coverage ≥ 50% for both sequences.

**Transcriptome analysis**

RNA samples were extracted from inoculated roots at post-inoculation 6 h and 3 dpi. Total RNA was extracted using a NucleoSpin RNA Plant (Macherey-Nagel). RNA samples (1µg each) were then sent to Macrogen for library preparation and subsequent sequencing. The generated libraries were sequenced by HiSeq-illumina (approximately 20 million reads per sample, paired-end). For plant transcriptome analysis, Tophat2 with the default setting (Kim *et al*., 2013) was used for sequence mapping to the reference *Arabidopsis thaliana* genome (TAIR10). The generated BAM files from the Tophat2 platform were used to analyze differentially expressed genes by cuffdiff with default settings (Trapnell *et al*., 2012). The following statistical analysis and visualization of data were conducted by R Bioconductor packages such as Cummerbund and ggplot2 (Ginestet 2011; Trapnell *et al*., 2012).

RNA-seq read sets obtained from CgE, CgP, and co-inoculated samples were subjected to adapter removal and quality filtering using Platanus_trim (version 1.0.7) with default parameters. The trimmed reads were classified using two sequential rounds of mapping. First, the trimmed reads were mapped to the *Arabidopsis thaliana* genome using HISAT (version 2.1.0) (Kim *et al*., 2015) with default parameters. Reads that were mapped onto the *Arabidopsis thaliana* genome were classified as originating from *Arabidopsis thaliana*. Next, we performed the second classification by mapping the reads that remained unmapped in the first classification onto CgE and CgP genomes. Reads that uniquely mapped to either CgE or CgP genomes were classified as originating from CgE or CgP, respectively. In addition, reads that could not be mapped to both genomes were classified as unmapped reads. Finally, reads that mapped to both genomes during the second step were further classified. The number of mismatches in the alignment reads were identified by the "XM" flag in the SAM output files and then the number of mismatches in the alignment against CgE and CgP genomes were compared. Based on the comparison results, reads were classified into CgE, CgP, and classified reads. The resulting CgE-RNA-seq derived CgE reads (CgE-CgE reads), CgP-RNA-seq derived CgP reads (CgP-CgP reads), co-inoculation-RNA-seq derived CgE reads (coinoc-CgE reads), and co-inoculation-RNA-seq derived CgP reads (coinoc-CgP reads) were used for differential expression analysis.

CgE-CgE reads and coinoc-CgE reads were mapped to the annotated genome of CgE; whereas, CgP-CgP reads and coinoc-CgP reads were mapped to the annotated genome of CgP using HISAT2 (version 2.1.0). The reads of each sample were used for transcript assembly and quantification of each transcript using the StringTie (version 1.3.4d) (Pertea *et al*., 2015) and

Ballgown R packages (Frazee *et al*., 2015). To identify differentially expressed genes, the edgeR package (Robinson *et al*., 2010) was used to compute the p-value and fold change. The p-value was used to identify genes expressed differentially between the paired datasets. Significantly, differentially expressed genes were identified using a FDR threshold of $\leq 0.05$ and a minimum two-fold change. RNAseq sequences used in this study are deposited in DDBJ (DRA009854, CgE=CfE).

#### Supplementary note

##### The expression of cell wall degrading enzymes (CWDEs)

GO related to carbohydrate metabolic process also dominated in CgE up-regulated genes following CgP co-inoculation (GO:0005975, FDR: 1.3E-8, Fig.6C). This promoted us to investigate the expression profiles of fungal CWDEs during the competition. Of the CgE 417 genes annotated for CWDEs, 98 and 8 genes were up- and down-regulated following CgP co-inoculation, respectively (Supplementary Table 9). Genes encoding degrading enzymes for pectin, hemicellulose, cellulose were especially enriched as up-regulated genes. These suggest that CgE activates an ability to degrade several types of plant cell wall components during the competition.

Of 417 CWDEs genes in CgP, 38 and 41 genes were up- and down-regulated, respectively, following CgE co-inoculation. Genes encoding degrading enzymes for pectin, hemicellulose, cellulose were especially enriched as up-regulated genes. Interestingly, three genes encoding chitin-binding domain (CBM18 and CBM50 modules) were also up-regulated during the competition (Supplementary Table 10).

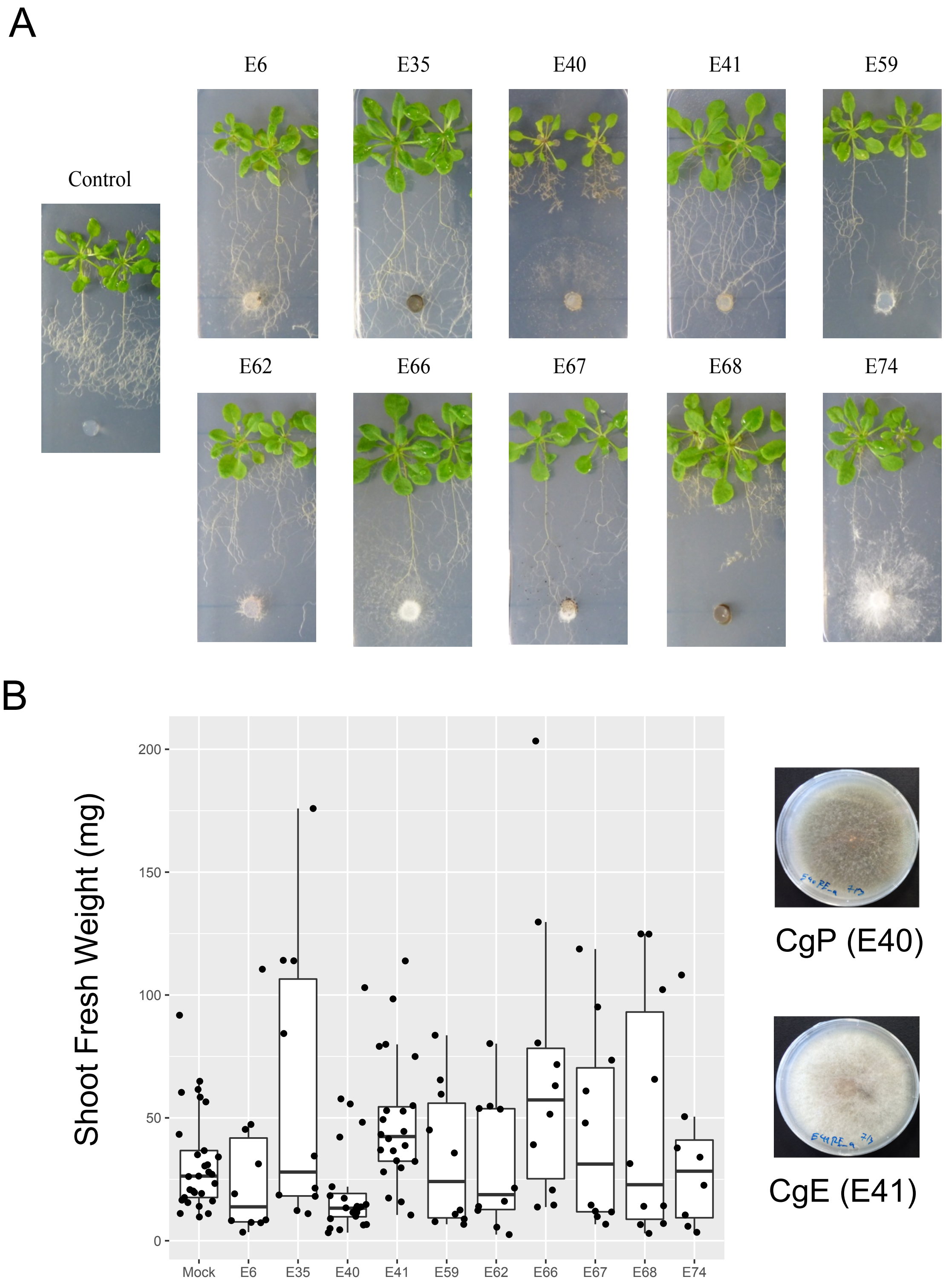


**Supplementary Fig. 1. Plant inoculation assay with the isolated fungal strains**
**(A)** Representative pictures of the inoculated *A. thaliana* plants by hyphae of 10 selected fungal isolates at 21 dpi. **(B)** Measurement of shoot fresh weight of *Arabidopsis* inoculated with the candidate fungi at 21 dpi. Each sample comprised at least 6 shoots. Each dot represents individual plant samples. The pictures of CgP (E40) and CgE (E41) grown in Mathur's media were shown (Right).

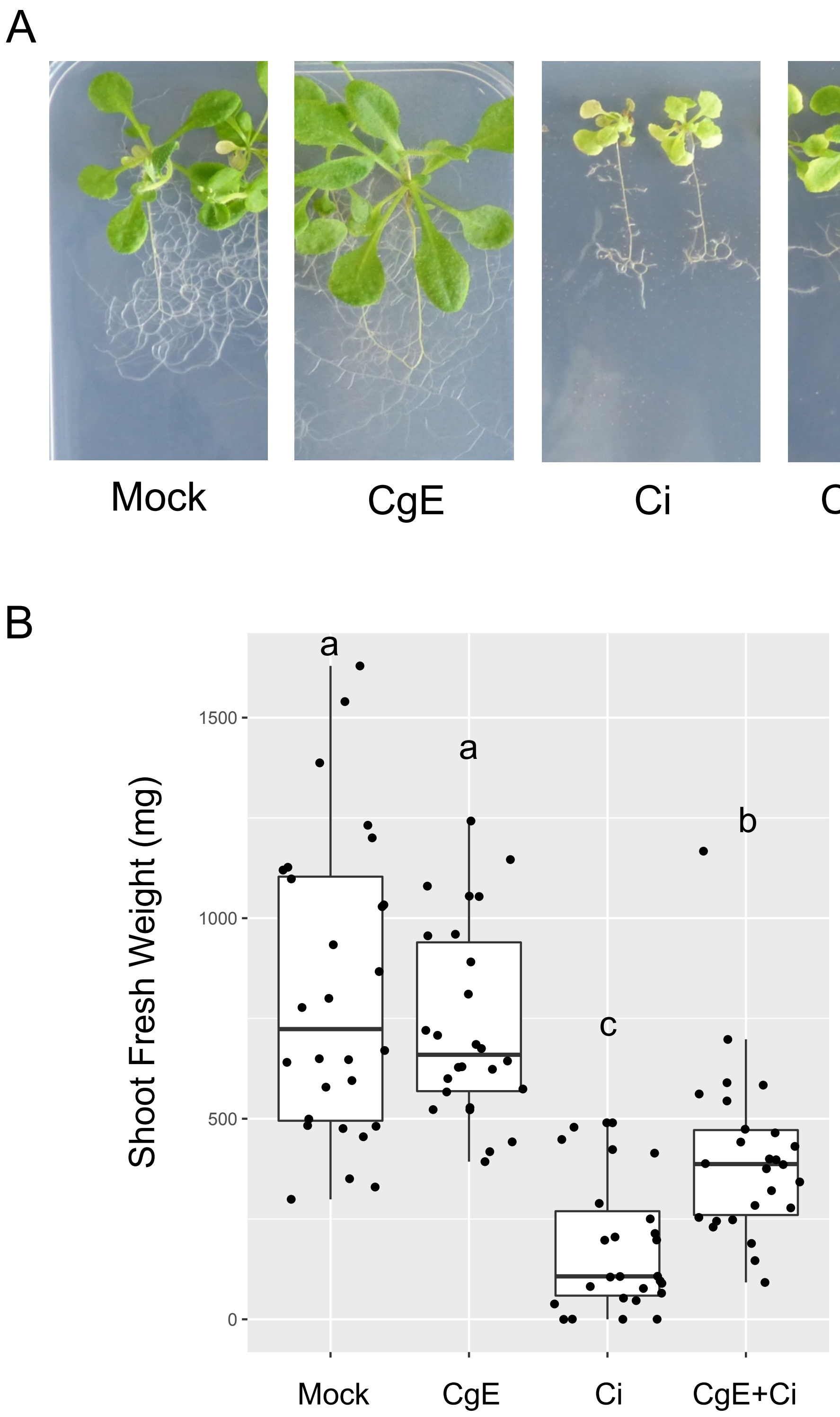


**Supplementary Fig. 2. CgE-mediated plant protection against pathogenic *Colletotrichum incanum***

**(A)** Morphology of plants treated with water, spores of endophytic *Colletotrichum* spp. (CgE), spores of pathogenic *C. incanum* (Ci) or co-inoculation of spores of pathogenic *C. incanum* (Ci) with those of endophytic *Colletotrichum* spp. (CgE) at 21 dpi on 1/2 MS agar media. **(B)** Shoot fresh weight of *Arabidopsis* from co-inoculation of pathogenic *Ci* with endophytic CgE assay at 21 dpi. The boxplot shows combined data from three independent experiments. Different letters indicate significantly different statistical groups (Tukey's HSD, $p < 0.05$).

#### Assembly Statistics

| | *C. orbiculare* | *C. fructicola* Nara gc−5 | CgE | CgP |
|---|---|---|---|---|
| Assembly size (Mb) | 88.3 | 55.6 | 64.5 | 57.7 |
| Number of scaffolds | 525 | 1241 | 35 | 21 |
| Median length (N50) | 428.89 kb | 112.81 kb | 4923.73 kb | 4922.22 kb |
| Gaps | 4761 | 4094 | 0 | 0 |

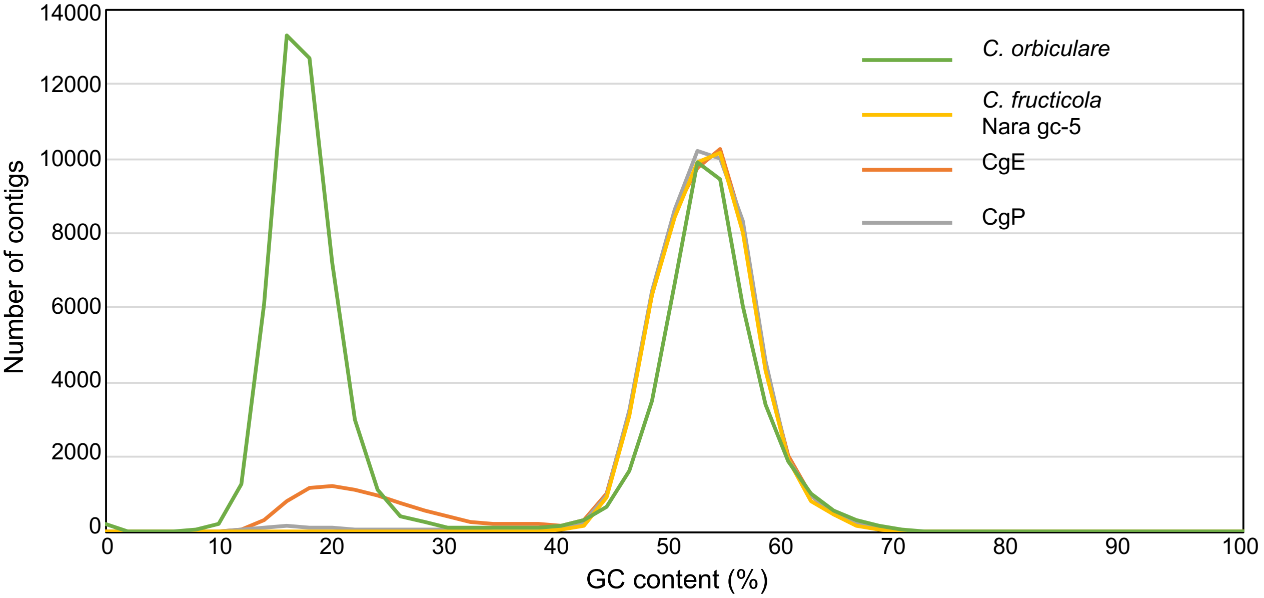


**Supplementary Fig. 3. Summary of genome assembly obtained by PacBio- and Illumina-derived reads.**
Assembly statistics of the newly obtained CgE and CgP genomes are shown in the upper table, together with those of previously published *C. orbiculare* and *C. fructicola* Nara gc-5 genomes. Compared with the previously published genomes, Lower number of scaffolds, longer N50 and no gaps indicate the high quality of CgE and CgP genomes. GC contents in 4 different *Colletotrichum* genomes are described in graph below. Genomes of *C. orbiculare* (Green) and CgE (Orange) have low GC content blocks (AT rich blocks).

A

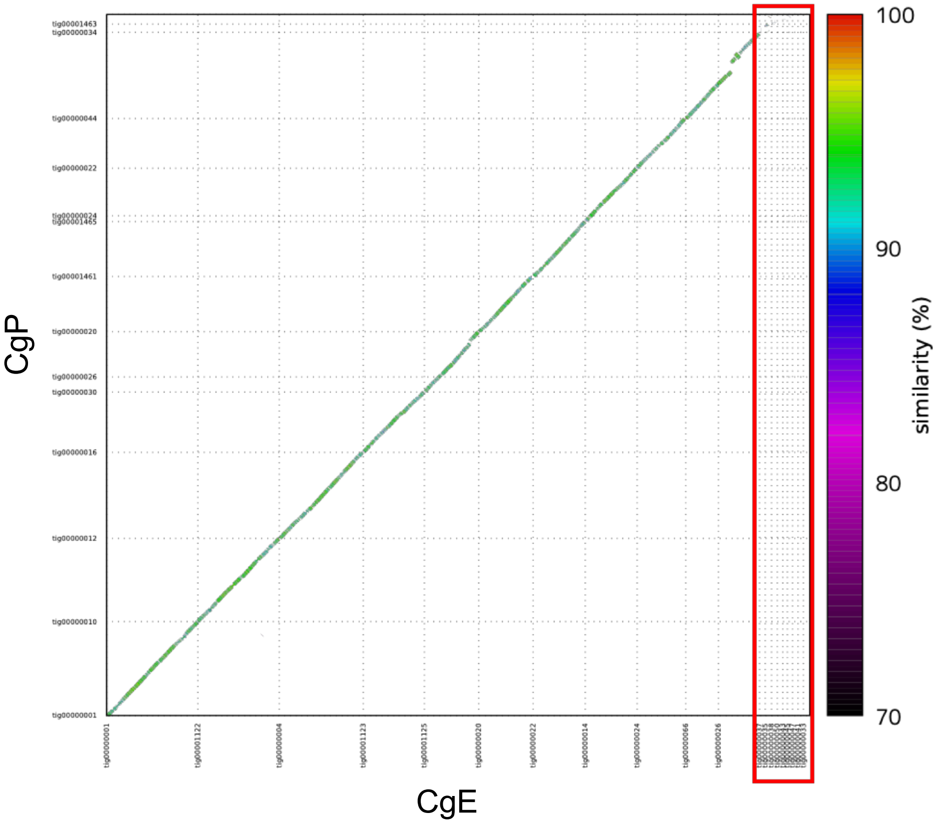


| scaffold | AT_rate (%) |
|---|---|
| tig00000001 | 10.48 |
| tig00001122 | 10.30 |
| tig00000004 | 10.42 |
| tig00001123 | 15.11 |
| tig00001125 | 10.09 |
| tig00000020 | 11.59 |
| tig00000022 | 9.89 |
| tig00000014 | 11.93 |
| tig00000024 | 12.91 |
| tig00000066 | 8.62 |
| tig00000026 | 10.01 |
| tig00000037 | 41.71 |
| tig00000035 | 35.41 |
| tig00000038 | 39.20 |
| tig00000050 | 32.27 |
| tig00000043 | 29.90 |
| tig00000045 | 29.09 |
| tig00000047 | 39.88 |
| tig00000041 | 26.26 |
| tig00000033 | 30.52 |

B

| | CgE | | CgP | | *C. fructicola* (Nara gc5) | | *C. orbiculare* | |
|---|---|---|---|---|---|---|---|---|
| | AT block | GC block | AT block | GC block | AT block | GC block | AT block | GC block |
| Total size (Mbp) | 8.40 | 56.09 | 1.14 | 56.54 | 0.21 | 55.39 | 45.34 | 44.75 |
| Average size (bp) | 14,586 | 98,927 | 4,112 | 204,840 | 914 | 44,599 | 11,015 | 52,029 |
| Median size (bp) | 8,000 | 56,000 | 2,000 | 35,000 | 1,000 | 19,618 | 785 | 16,000 |
| GC content (%) | 23.87 | 53.69 | 25.27 | 53.71 | 33.73 | 53.46 | 18.10 | 55.01 |
| Number of genes | 13 | 15,999 | 3 | 15,081 | 5 | 15,376 | 165 | 13,182 |
| Gene density (gene/Mb) | 1.55 | 285.23 | 2.64 | 266.75 | 23.26 | 277.58 | 3.64 | 294.60 |

**Supplementary Fig. 4. Alignment between CgE and CgP genomes.**
**(A)** Dot plot was obtained using Nucmer and a modified version of Mummerplot in the Mummer package. Color code represents similarity (%) between CgE and CgP genomes (from red (high) to black (low)). Roughly, both genomes show ~ 90% similarity each other. Several gaps represented by white color on the diagonal line were detected between CgE and CgP genomes (For the detail, see also Supplementary Fig.5). Genomic regions highlighted by red square represent AT rich regions (GC contents < 40 %). **(B)** Summary of AT rich block and GC rich block among different *Colletotrichum* strains including CgE and CgP. The number of genes at AT rich blocks is also described.

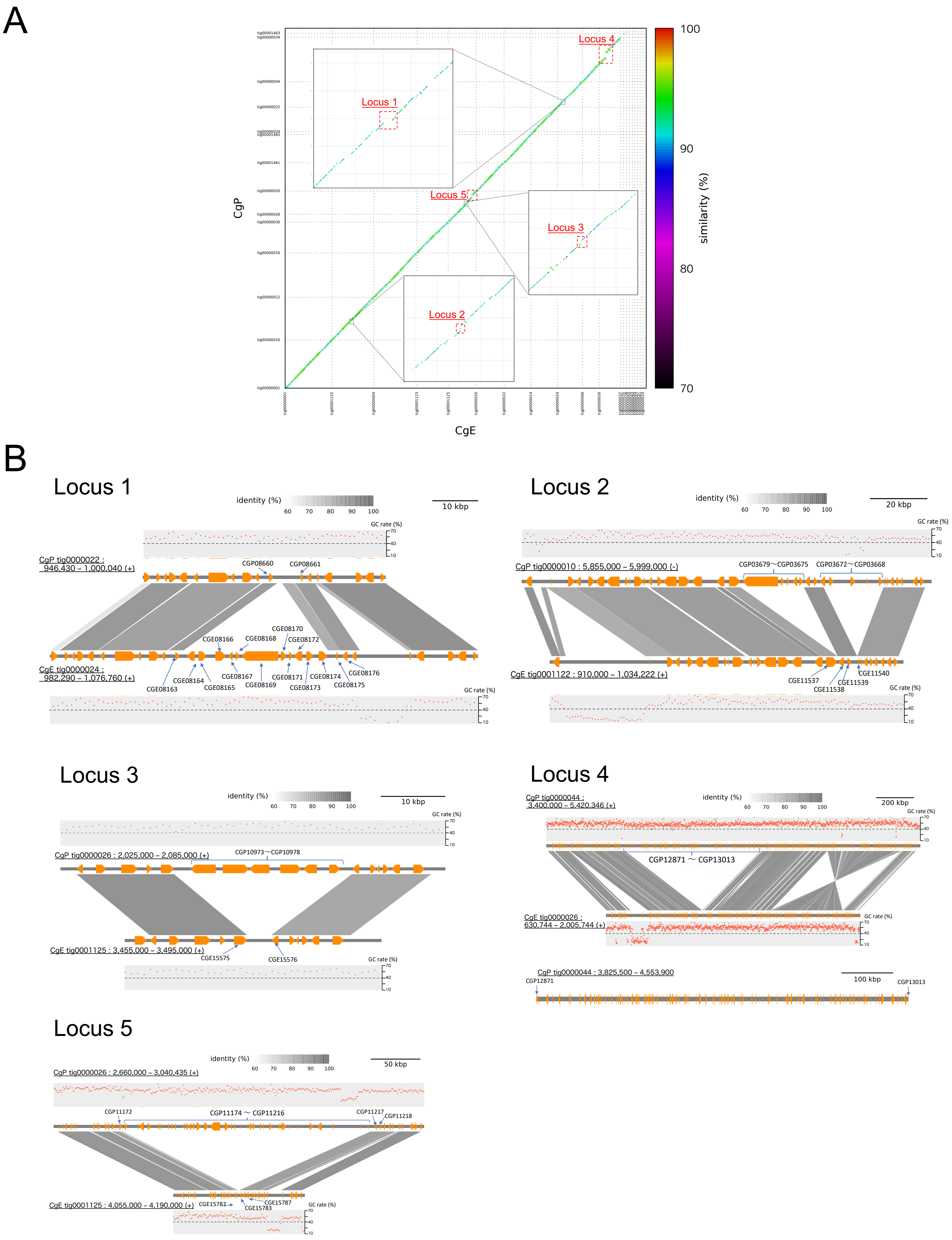


**Supplementary Fig. 5. Genomic locus and genes specific either to CgE or CgP (A, B)** Five genomic locus specific to either in CgE or CgP GC rich blocks were extracted from the genomic alignment plot as shown in (A). CgE or CgP specific genes (CgE or CgP genes not shared by the other) located in each specific locus were identified by reciprocal BLAST analysis and dot plot in (A). Genes were represented by orange polygons. Vertical bars connecting adjacent genomic structures indicate BLAST hit blocks in the comparison between the two adjacent genomic scaffolds (B). GC rate indicates GC content per 1 kb window.

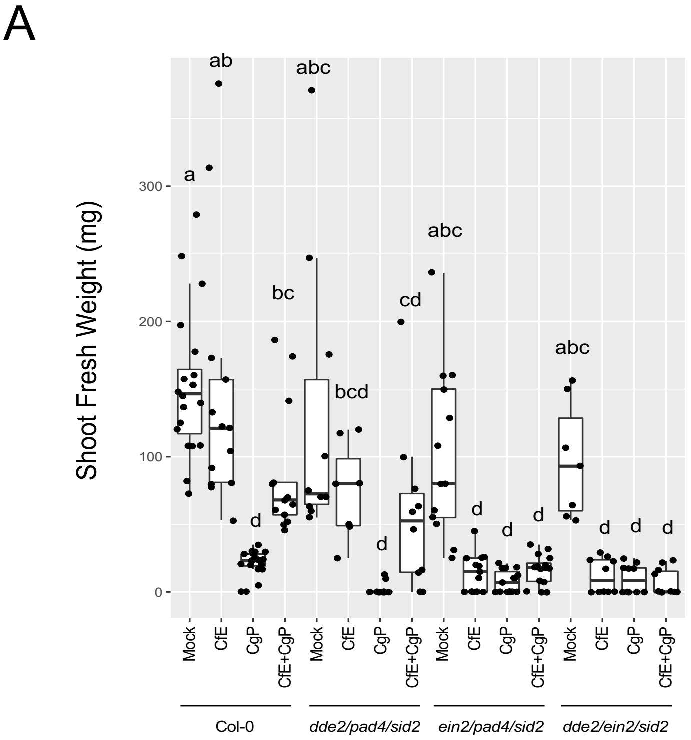


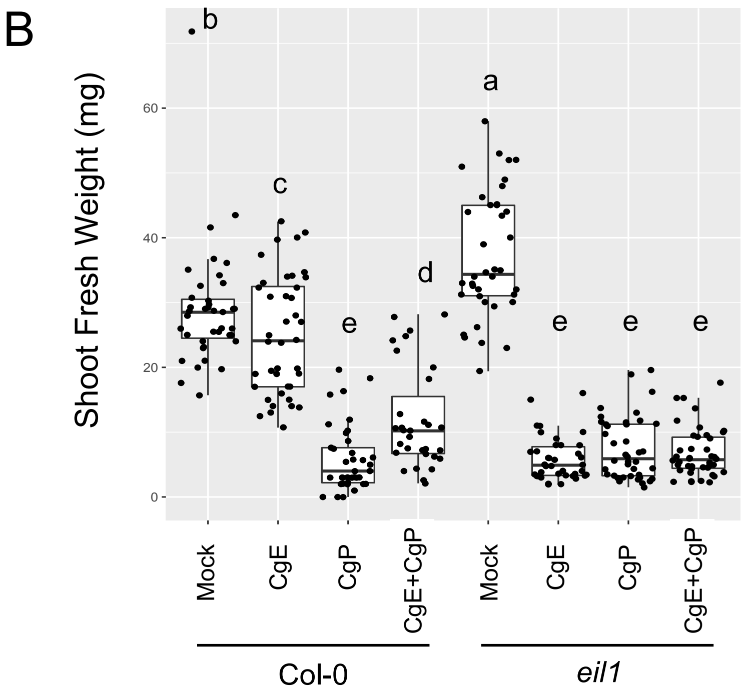


**Supplementary Fig. 6. Requirement of ethylene signaling for the endophytic lifestyles of CgE and its host protection**

**(A)** Measurement of the shoot fresh weight of *Arabidopsis* hormonal triple mutants in the co-inoculation assay at 14 dpi**.** The boxplot shows combined data from three independent experiments. Different letters indicate significantly different statistical groups (Tukey's HSD, $p$ < 0.05). *A. thaliana* mutants with *ein2* mutation show defective in the endophytic lifestyles of CgE and its host protection. **(B)** Measurement of shoot fresh weight of wild-type *Arabidopsis* and ET-related *eil1* mutants in the co-inoculation assay at 14 dpi. Each sample comprised around 20 shoots per experiment. The boxplot shows combined data from two independent experiments. Different letters indicate significantly different statistical groups (Tukey's HSD, $p$ < 0.05).

A

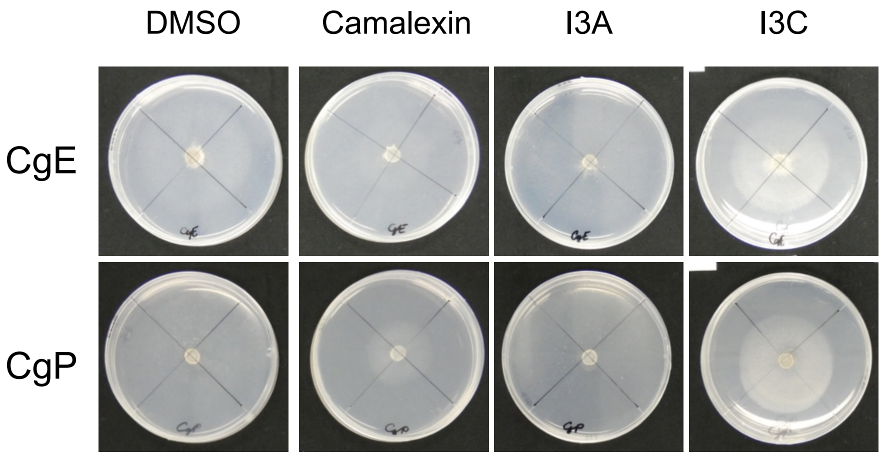


B

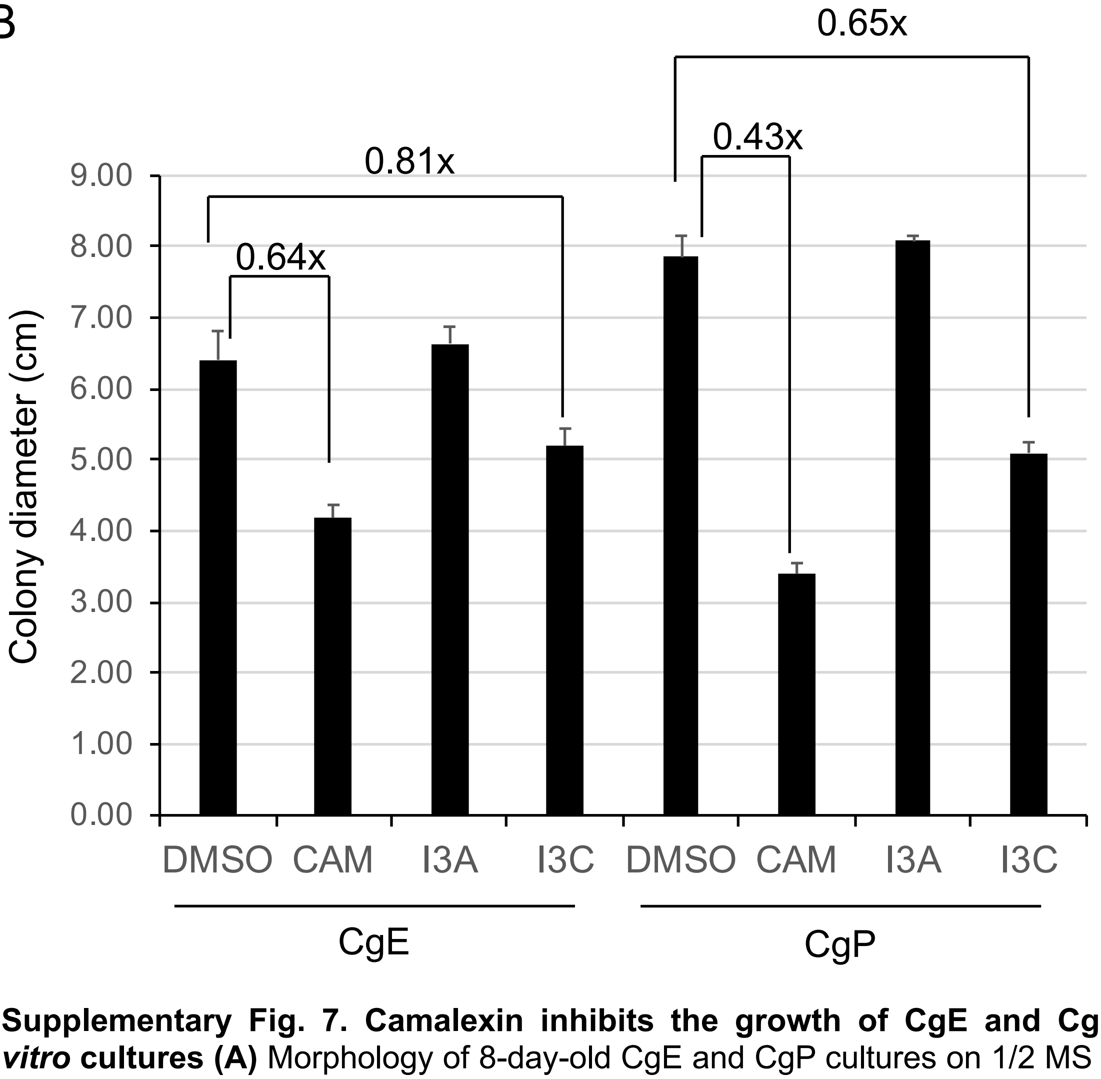


**Supplementary Fig. 7. Camalexin inhibits the growth of CgE and CgP *in vitro* cultures (A)** Morphology of 8-day-old CgE and CgP cultures on 1/2 MS agar media supplemented with Trp-derived secondary metabolites [camalexin (Cam), indole-3-ylmethylamine (I3A), indole-3-carbinol (I3C), and 4-hydroxyindole-3-carbonitrile (4OHICN)]. **(B)** Colony diameter of 8-day-old CgE and CgP colonies cultured on 1/2 MS agar media supplemented with tryptophan-derived secondary metabolites. Data are presented as the mean ± SD of at least 4 different colonies. x means ratio of Chemical/DMSO. The ratio of CAM/DMSO in CgE is higher than that in CgP.

Supplementary Fig.8

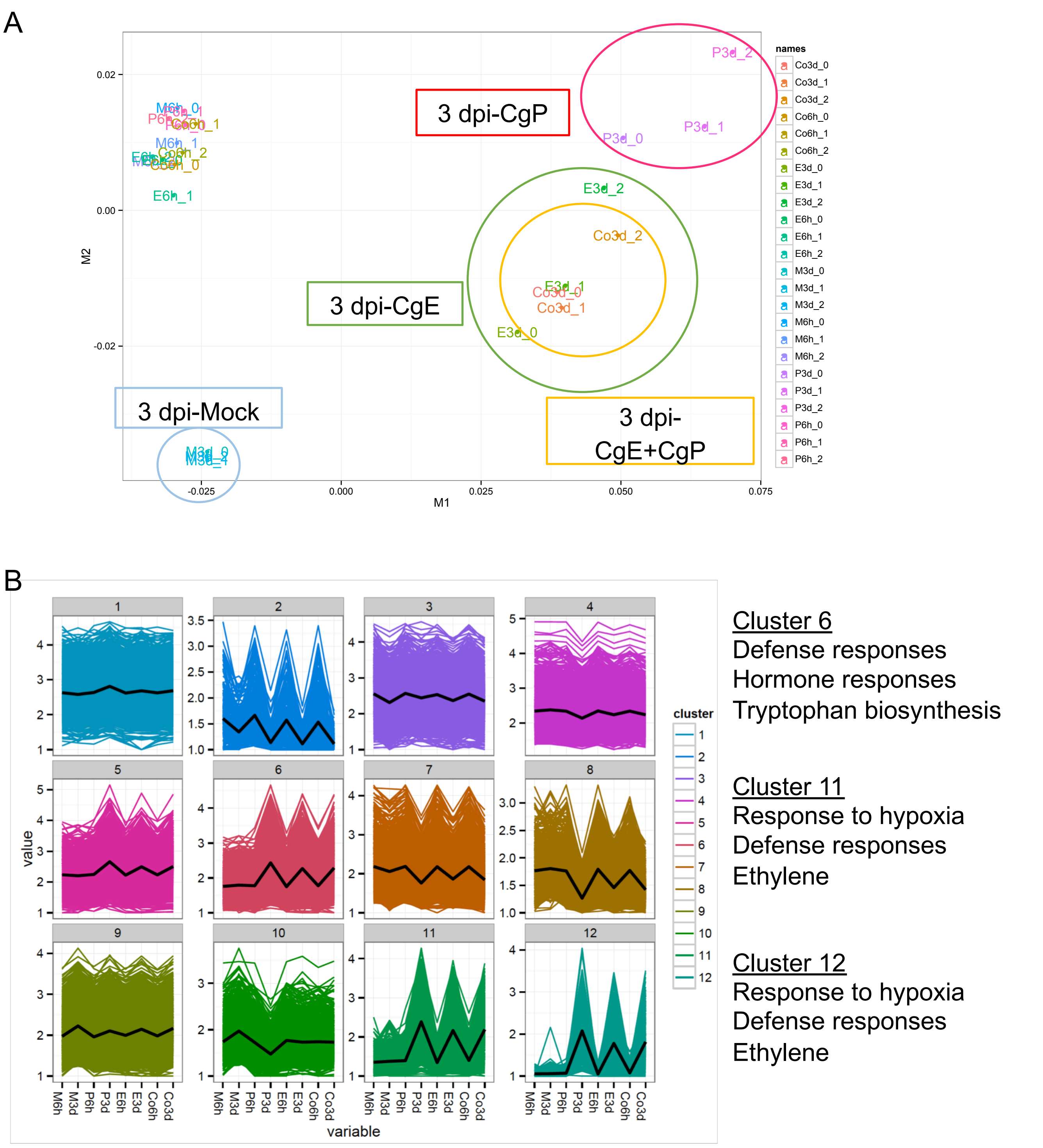


**Supplementary Fig. 8. Plant-side transcriptome analysis during the root colonization of CgE and CgP**

**(A)** Multidimensional analysis chart generated from plant transcriptome profile during association with CgE and/or CgP at post-inoculation 6 h and 3 dpi. M = Mock, E = CgE, P = CgP, and Co = CgE + CgP. The circles are drawn as they cover the position of each replicate. **(B)** The expression patterns of plant genes differentially expressed in a pairwise comparison (FDR < 0.01) –cluster analysis divided by k-mean (K = 12). Genes listed in each cluster were subjected to Go analysis.

### Supplementary Fig.9

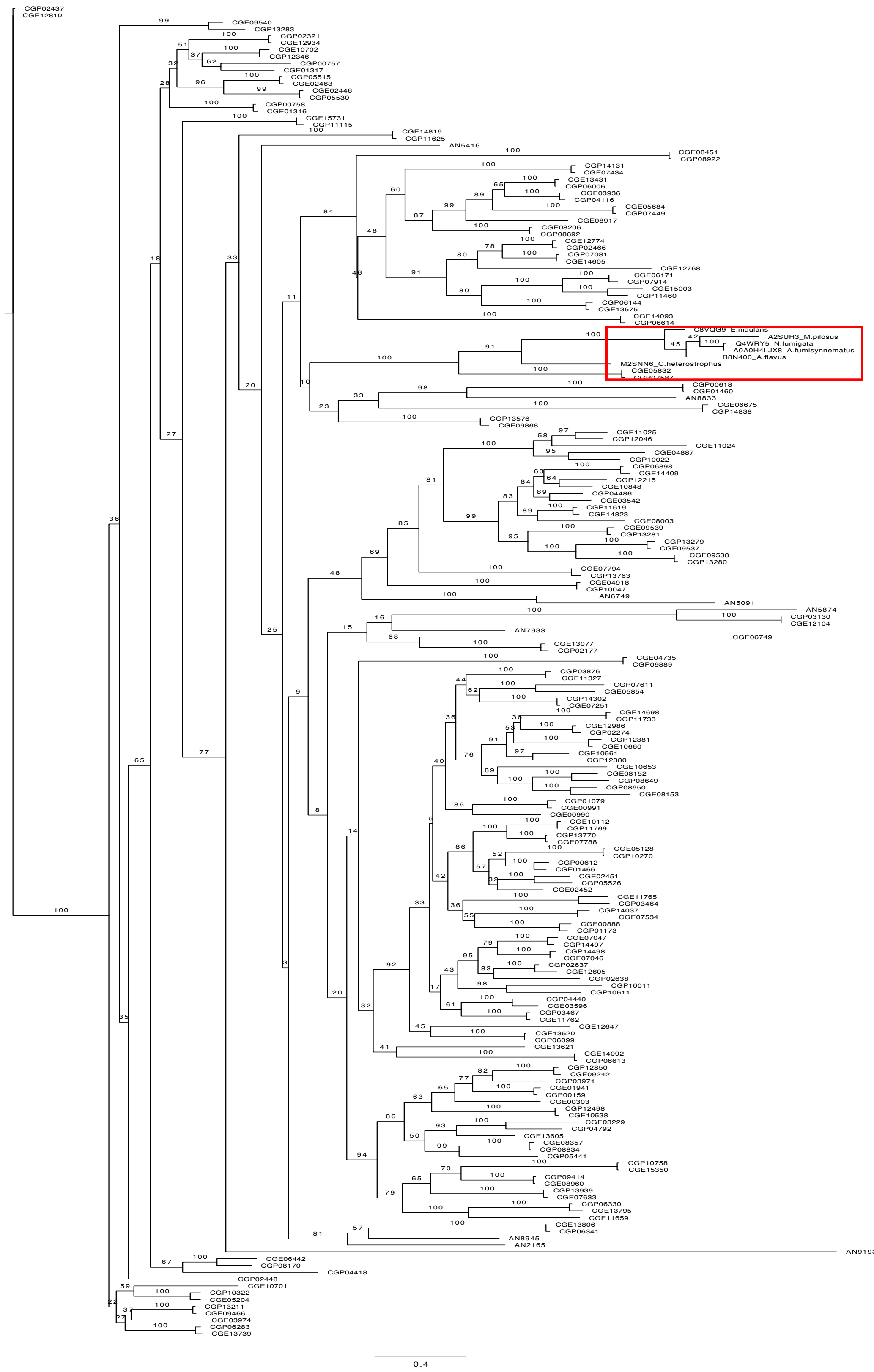

**Supplementary Fig. 9. Phylogenic tree of LaeA-like methyltransferases in CgE and CgP**

Phylogeny of 94 CgE and 86 CgP LaeA-like methyltransferases (LaeA and llm) are described. The phylogenic tree was generated by RaxML. The red square represents LaeA in other fungi (C8VQG9, A2SUH3, Q4WRY5, A0A0H4LJX8, B8N406, and M2SNN6) and the corresponding CgE and CgP genes (*CGE05832* and *CGP07587*). llm in *A. nidulans* were also described in the tree (AN2165, AN8945, AN7933, AN5091, AN6749, AN8833, AN5416, AN5874, and AN9193).

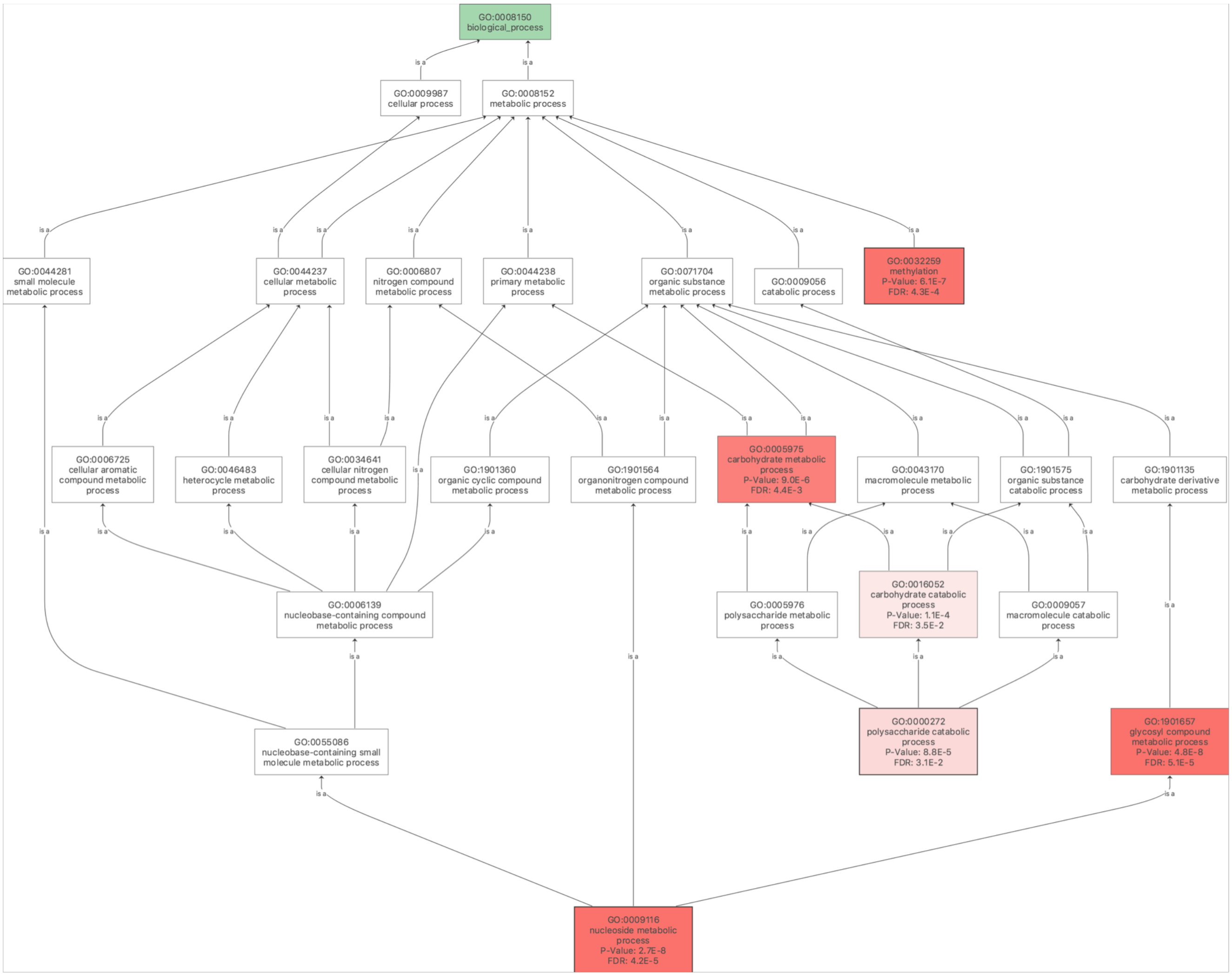


**Supplementary Fig. 10. Enriched GO terms in 564 CgP genes up-regulated in co-inoculated samples compared with it alone.**

564 CgP genes significantly up-regulated in co-inoculated samples compared with its alone were subjected to GO analysis ($\log_2$ FC >1, FDR < 0.05). This picture shows the result of Go term analysis related to biological process. Especially, Go terms such as methylation, glycosyl compound metabolic process, and nucleoside metabolic process were significantly enriched. GO analysis was conducted by Blast2go with default setting.

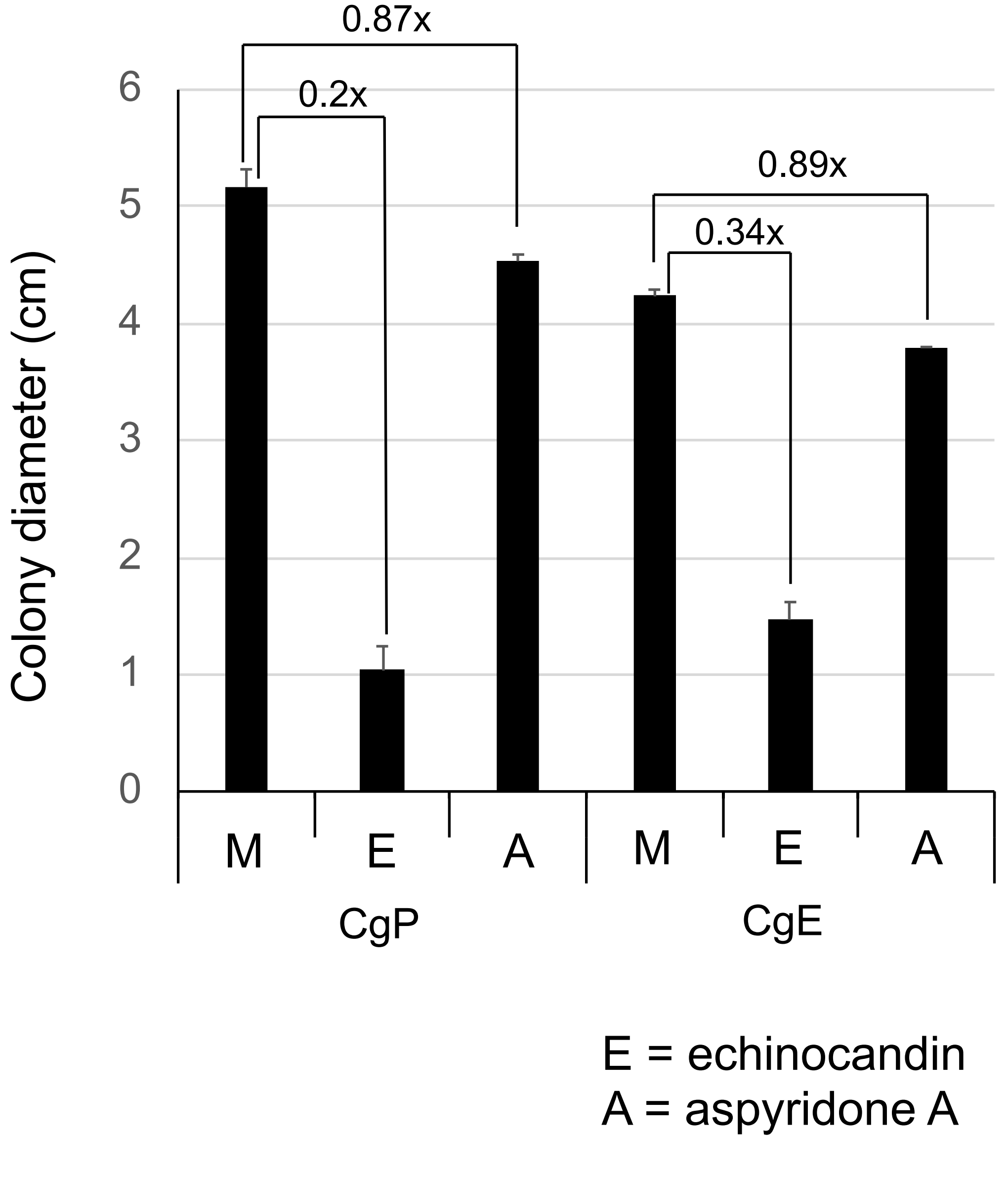


**Supplementary Fig. 11. Fungal toxin assay *in vitro* cultures**
Colony diameter of 8-day-old CgE and CgP colonies cultured on PDA media supplemented with or without fungal toxin echinocandin B (4 μg/ml, E) or aspyridone A (4 μg/ml, A). Data are presented as the mean ± SD of at least 4 different colonies. x means ratio of Fungal toxin/Mock. The ratio of echinocandinB/Mock in CgE is higher than that in CgP.
